# Efficient *in vivo* mammalian neuron editing using peptide-mediated CRISPR enzyme delivery

**DOI:** 10.1101/2025.11.26.690638

**Authors:** Brigette D. Manohar, Meika Travis, Vikas Munjal, Christopher M. Baehr, Lily M. A. Rahnama, Min Hyung Kang, Rania F. Haddad, Kunica Asija, Eric A. Noel, Navya Peddireddy, Runvir S. Chouhan, Rohit Sharma, Stacia K. Wyman, Netravathi Krishnappa, Deirdre A. Killebrew, Song Hua Li, Kathy J. Snow, Addison K. Byrne, Chandra S. Boosani, P. Anthony Otero, John Bringas, Allison A. O’Brien, Matthew T. Rocco, Madison P. Zimmerly, Isabelle Gray, Gunita K. Sran, Mia I. Patel, Emily F. Seidensticker, Ethan Saville, Yaned Gaitan, Amanda L. Schmelzle, Pradeep Nag B. Subramanyam, Lluis Samaranch, Jonathan A. Green, Kevin D. Wells, Alexander J. Ehrenberg, Stephen A. Murray, Claire D. Clelland, Niren Murthy, Russell R. Lonser, Piotr Hadaczek, Victor S. Van Laar, Krystof S. Bankiewicz, Ross C. Wilson

## Abstract

CRISPR-mediated genome editing of the central nervous system (CNS) has the potential to revolutionize the treatment of neurological disorders, including neurodegenerative disorders such as Huntington’s disease (HD). However, the development of CRISPR therapeutics for the CNS has been hindered by challenges associated with delivery, specifically the lack of a clinically compatible, non-viral delivery technology facilitating genome editing of neurons *in vivo*. For most indications, two key obstacles must be overcome before therapeutic genome editing of the brain is feasible: non-toxic intracellular delivery of CRISPR cargo into neurons and establishment of strategies enabling targeted brain regions to be edited efficiently. While viral vectors have shown promise in pre-clinical models, non-viral approaches present distinct advantages: ease of manufacture as well as the transient presence of CRISPR machinery, which tempers risks of genotoxicity and immunogenicity. Peptide-enabled ribonucleoprotein (RNP) delivery of CRISPR (PERC) has emerged as a promising non-viral delivery strategy for CRISPR enzymes with initial use in primary human immune cells. In this study, we report the development of Neuro-PERC, a streamlined and optimized approach for *in vivo* editing of mammalian neurons. Administration of Neuro-PERC reagents via convection-enhanced delivery (CED) mediated efficient and well-tolerated neuronal genome editing. Neuro-PERC enabled robust neuronal editing in the brain of both small and large animal reporter models, and increased survival in a severe murine model of Huntington’s disease. These results establish CED-administered Neuro-PERC as a candidate delivery technology to hasten clinical translation of CRISPR-based therapies for diseases of the CNS.

**Summary:** Neuro-PERC, a peptide-mediated CRISPR enzyme delivery technology, enables efficient in vivo mammalian neuronal editing in the brain of mice and pigs, extending survival in a murine model of Huntington’s disease when administered via convection-enhanced delivery (CED).

## INTRODUCTION

CRISPR-based genome editing enables prevention or treatment of disease by correcting mutations, modulating gene expression, or introducing therapeutic DNA (*1*). While *ex vivo* delivery of CRISPR-Cas9 followed by subsequent cell transplantation rapidly showed clinical success (*2*), the full potential of therapeutic genome editing hinges on the ability to administer genome editing reagents directly into the body (*3*). Candidate genome editing therapies for CNS diseases are progressing towards clinical use (*4*), but there is a dearth of safe and effective *in vivo* delivery technologies for genome editing effectors in the brain (*5*). Overcoming this limitation could enable transformative treatments for genetic neurological disorders of the CNS such as Huntington’s disease (HD) (*6*), that currently lack disease-modifying therapies (*7, 8*).

Despite proof-of-concept success in animal models, adenoviral-associated virus (AAV)-based CRISPR delivery has shown limited clinical benefit to-date. Subretinal delivery of a AAV5-based CRISPR-Cas9 therapy for LCA10 retinal disease yielded marginal improvement (*9*), and high-dose systemic AAV9-based CRISPR delivery in a Duchenne muscular dystrophy trial caused a fatal immune response (*10*). Although AAV-based platforms can efficiently target the brain in animal models (*11, 12*), they bear CRISPR-specific safety concerns, including frequent genomic integration at double-stranded breaks, prolonged effector expression that increases off-target editing rates, and immune responses to the encoded microbial CRISPR protein (*13*).

Non-viral platforms enact transient delivery and may reduce genotoxicity and immunogenicity (*14*), but to date, have generally shown lower editing efficiencies and more limited brain distribution than AAV (*15–17*). These trade-offs highlight the need for a delivery strategy that combines the safety of non-viral systems with the efficiency and tissue dispersion of AAV. An optimal *in vivo* brain delivery platform for CRISPR would feature transient effector presence to limit off-target editing and immunogenicity, absence of DNA cargo (eliminating risks of integration), efficient intracellular delivery, small particle size for broad tissue coverage, scalable manufacturing, and robustness to sterile filtration and freeze/thaw cycles.

Delivery of CRISPR enzymes in active ribonucleoprotein (RNP) form satisfies several of these criteria. CRISPR RNP is inherently transient following intracellular delivery, with a ∼24 h half-life (*18*), and has a small hydrodynamic diameter (≤ 20 nm) (*19, 20*) that should support efficient diffusion through brain parenchyma (*21*), akin to ∼25 nm AAV virions (*17, 22*). In contrast, lipid nanoparticles that carry CRISPR mRNA (*23, 24*) typically exhibit > 100 nm diameters, exceeding the estimated 35–64 nm size cutoff for effective diffusion in the brain (*25*). Despite the appeal of CRISPR delivery in RNP format, enzymes such as CRISPR-Cas9 remain limited in their ability to mediate high-efficiency and widespread genome editing *in vivo*.

Peptide-enabled RNP delivery of CRISPR (*e.g.* PERC) has shown promise in tissue culture (*26–30*) and *in vivo* (*31*), but peptide:RNP complexes typically form > 60 nm aggregates anticipated to distribute poorly in brain parenchyma (*27*). We hypothesized that PERC could be adapted for *in vivo* brain delivery by establishing a monodisperse RNP formulation compatible with convection -enhanced delivery (CED), a clinically validated administration route that supports broad dispersion of macromolecules in brain tissue and can promote uptake into cells via the pressurized infusion (*32–34*). We report Neuro-PERC, a monodisperse CRISPR RNP–peptide complex optimized for neuronal editing via CED. *In vitro*, biophysical, and *in vivo* studies in mice and pigs show that Neuro-PERC enables efficient, non-toxic, minimally immunogenic neuronal editing and extends lifespan in an aggressive murine HD model, highlighting its potential as a therapeutic platform for *in vivo* brain genome editing.

## RESULTS

### Optimized Neuro-PERC formulation mediates efficient genome editing in neural cells

Our previous work established PERC in primary human immune cells (*27*), and a complementary report established peptide-assisted genome editing (PAGE) (*28*); both studies demonstrated efficient delivery by pairing engineered nuclear localization signal (NLS)-rich *S. pyogenes* Cas9 (henceforth Cas9 unless another species of origin is noted) enzymes with engineered amphiphilic delivery peptides that combine the canonical cell-penetrating peptide TAT (*35*) covalently linked to an endosomolytic sequence derived from influenza HA2 (*36*). To evaluate the PERC for mammalian neurons *in vivo*, we first tested the *in vitro* editing capabilities in primary neural progenitor cells (NPCs) derived from Ai9 reporter mice (*37*), which express tdTomato red fluorescent protein (RFP) upon excision of a ∼750 bp polyA transcription repressor sequence using a pair of Cas9 guide RNAs (gRNAs) (*38*). Employing a Cas9 construct bearing an optimized three-NLS configuration (*39*) that enhances PERC efficiency (*27*), NPCs were treated with various peptide:RNP formulations added directly to the media to assess genome editing via flow cytometry (Fig. 1A).

**Fig. 1:**
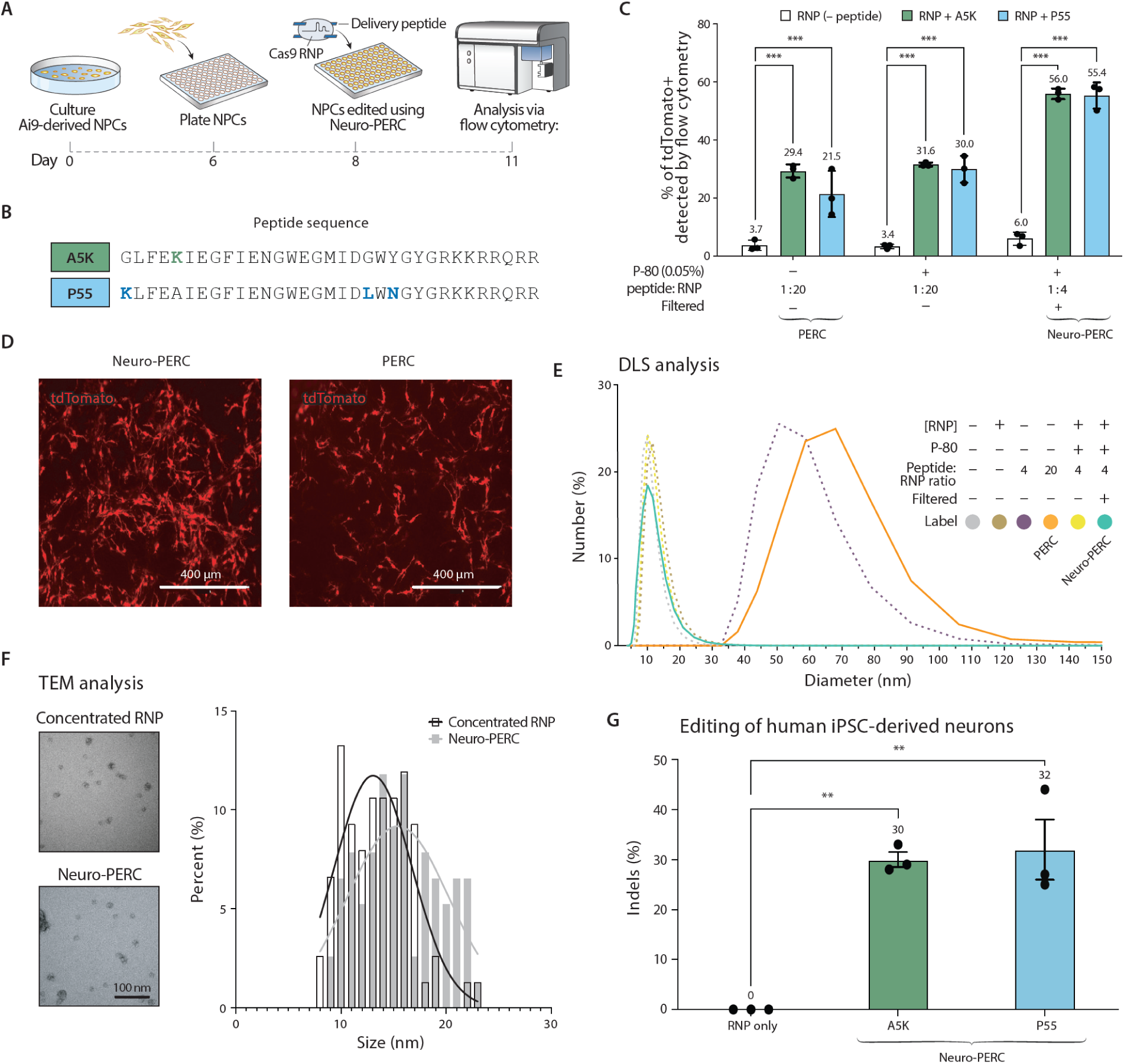
Optimization and characterization of Neuro-PERC for neuronal genome editing *in vitro*. **(A)** Schematic workflow for genome editing in Ai9-derived NPCs. NPCs were plated as a monolayer on day 6 of culture, followed by direct addition of Cas9 RNP and delivery peptides to the culture medium on day 8. Genome editing efficiency was quantified by flow cytometric detection of tdTomato⁺ fluorescence. **(B)** Amino acid sequences of delivery peptides A5K and P55, with differing residues shown in bold and color-coded (blue: P55, green: A5K). **(C)** Quantification of tdTomato⁺ NPCs following treatment with Cas9 RNP alone or formulations containing A5K or P55 via flow cytometry. Experimental variables included peptide-to-RNP ratio, presence or absence of P-80, and formulation filtration. Data represent biological replicates (n = 3) and are shown as means ± SD. Statistical analysis was performed by two-way ANOVA. ***P < 0.001. **(D)** Representative microscopy of tdTomato^+^ NPCs from the corresponding experimental conditions. Scale bar, 400 µm. **(E)** DLS analysis showing particle-size distributions for indicated formulations. Curves represent formulations prepared with differing peptide:RNP ratios (4:1 or 20:1), with or without filtration or with or without 0.06% P-80. Color coding corresponds to formulations listed in the adjacent table. **(F)** TEM images of concentrated RNP-only and Neuro-PERC with corresponding size-distribution profiles. **(G)** Genome editing in iPSC-derived neurons treated with A5K- or P55-based Neuro-PERC targeting B2M. Both formulations achieved ∼30% indels, while RNP only showed negligible activity. Cells were washed 2 hours post-treatment and collected after 1 week for genomic analysis. Each data point represents one biological replicate (n = 3). Data are presented as mean ± SD and were analyzed by two-way

We tested peptide A5K, previously shown to enhance editing in primary human immune cells (*27*), as well as peptide P55, a modified variant incorporating three amino acid substitutions known to increase PERC activity (*29*) (Fig. 1B). Treatment with A5K or P55 using the previously reported formulation resulted in 29.4% ± 1.1% and 21.5% ± 4.2% tdTomato^+^ cells, respectively (Fig. 1C), compared with to 3.7 ± 0.9% tdTomato^+^ cells in RNP-only controls (*P* < 0.001 vs. control for both A5K and P55). Following the addition of PERC to NPCs, tdTomato^+^ cells exhibited typical neuronal morphology with no apparent alterations (Fig. 1D). In considering use of PERC for *in vivo* brain delivery, we suspected that the capacity for amphipathic delivery peptides to induce aggregation in the context of the previously reported formulation – featuring a 20:1 peptide:RNP ratio and no stabilizing buffer components – may limit diffusion through brain parenchyma (*25*). Analysis with dynamic light scattering (DLS) showed that the *ex vivo* formulation particle size (∼60 nm) (*27*) was substantially larger than the ∼13 nm Cas9 RNP (*27, 40*), a factor known to influence diffusion in dense brain tissue (*41*) (Fig. 1E, orange trace). We hypothesized that reducing the peptide:RNP ratio could yield smaller particles, yielding a formulation suitable for use in the brain. We evaluated various peptide:RNP ratios (4:1, 8:1, 10:1, 20:1) with peptide A5K. The 4:1 ratio achieved the highest tdTomato⁺ cell editing in mouse NPCs at a reduced dose of 50 pmol, with DLS showing a modest particle size decrease to ∼50 nm, which is still substantially larger than a single RNP particle and indicative of aggregation (Fig. 1E, purple trace; Fig. S1).

Polysorbate 80 (henceforth P-80, also known as Tween 80) is a non-ionic surfactant that has been shown to diminish nanoparticle agglomeration (*42*) and has been used in FDA-approved drug formulations (*43*). We next tested whether inclusion of P-80 could reduce the average particle size of the PERC formulation. With the addition of 0.06% P-80, we observed no detectable morphological differences in the mouse NPCs (Fig. 1D) and editing efficiencies of 31.6% ± 0.3% or 29.9% ± 2.3% tdTomato^+^ cells with peptide formulations (*P* < 0.001 vs. control for both A5K and P55) (Fig. 1C). Formulations containing low-dose P-80 yielded smaller, more monodisperse particles (Fig. 1E, yellow and cyan traces) resembling RNP alone (Fig. 1E, gray trace) without diminishing editing efficiency.

We next assessed compatibility of the formulation with sterile filtration, a critical consideration for *in vivo* administration (*44*). Using PAGE (polyacrylamide gel electrophoresis) to assess peptide retention after filtration through a 0.2 μm cellulose acetate filter, we found that PERC formulations lacking P-80 resulted in substantial peptide loss, an observation consistent with a scenario wherein some large peptide:RNP aggregates (as observed by DLS) exceed the pore size of the filter. In contrast, the PERC formulation optimized for *in vivo* use (4:1 peptide: RNP; P-80, filtered) – henceforth Neuro-PERC – could be filtered and retained most of the peptide (an average 85.8% ± 0.05% peptide retained), suggesting that the inclusion of P-80 improves formulation compatibility with sterilization protocols (Fig. S1). To evaluate the impact of filtration on editing efficiency, NPCs were dosed with Neuro-PERC. Flow cytometry showed robust editing: 55.9% ± 0.9% and 55.4% ± 2.2% tdTomato^+^ cells for A5K or P55, respectively, compared with 6.03 ± 1.1% tdTomato^+^ cells in RNP-only controls (*P* < 0.001 vs. control for both A5K and P55) (Fig. 1C).

Next, analysis via DLS and transmission electron microscopy (TEM) confirmed that Neuro-PERC consists of small, monodisperse particles (∼17 nm hydrodynamic diameter via DLS; ∼17 nm median diameter via TEM) (Fig. 1E, cyan trace; Fig. 1F), significantly smaller than the previously reported PERC formulations and consistent with monodisperse RNP particles. A freeze-thaw cycle had limited impact on particle size, as did prolonged incubation at room temperature (Fig. S1).

To further assess translational potential in a CNS-relevant tissue culture context, Neuro-PERC was tested in post-mitotic human iPSC-derived neurons (*45*) by targeting the *B2M* gene. Formulations with peptide A5K or P55 produced robust genome editing, yielding 30 ± 1.7% and 32 ± 6.9% indel formation, respectively (*P* < 0.01 vs. control for both A5K and P55) (Fig. 1G) but subsequently induced neuronal cell death (Fig. S1). Attaining even moderately efficient genome editing in post-mitotic human neurons *in vitro* is itself a notable observation considering the recalcitrant challenge this represents for the field (*46*), likely due to the fragility of neuron-like cells cultured in isolation. Neuro-PERC may prove even more useful in human *in vitro* human neuron models with context-specific optimization.

These results demonstrate the ability of optimized peptide:RNP formulations to function efficiently in both human and mouse cellular contexts. Neuro-PERC collectively embodies the properties of a CRISPR delivery technology with the potential to support *in vivo* use in diverse settings.

### Neuro-PERC mediates efficient, well-tolerated neuronal genome editing in the mouse brain

Convection-enhanced delivery (CED) enables widespread distribution of molecules in the brain and has been used to deliver Cas9, including self-delivering variants, for genome editing in mice (*38, 46, 47*) as well as AAV-based platforms (*47, 48*). We hypothesized that Neuro-PERC, combined with the broad distribution enabled by CED, would facilitate efficient neuronal editing in Ai9 mice (Fig. 2A). Stereotactic CED was performed unilaterally or bilaterally into the striatum at AP +0.8 mm, ML ±2.0 mm, DV −2.6 mm relative to bregma to enable widespread distribution within the striatal target region (Fig. 2B).

**Fig. 2:**
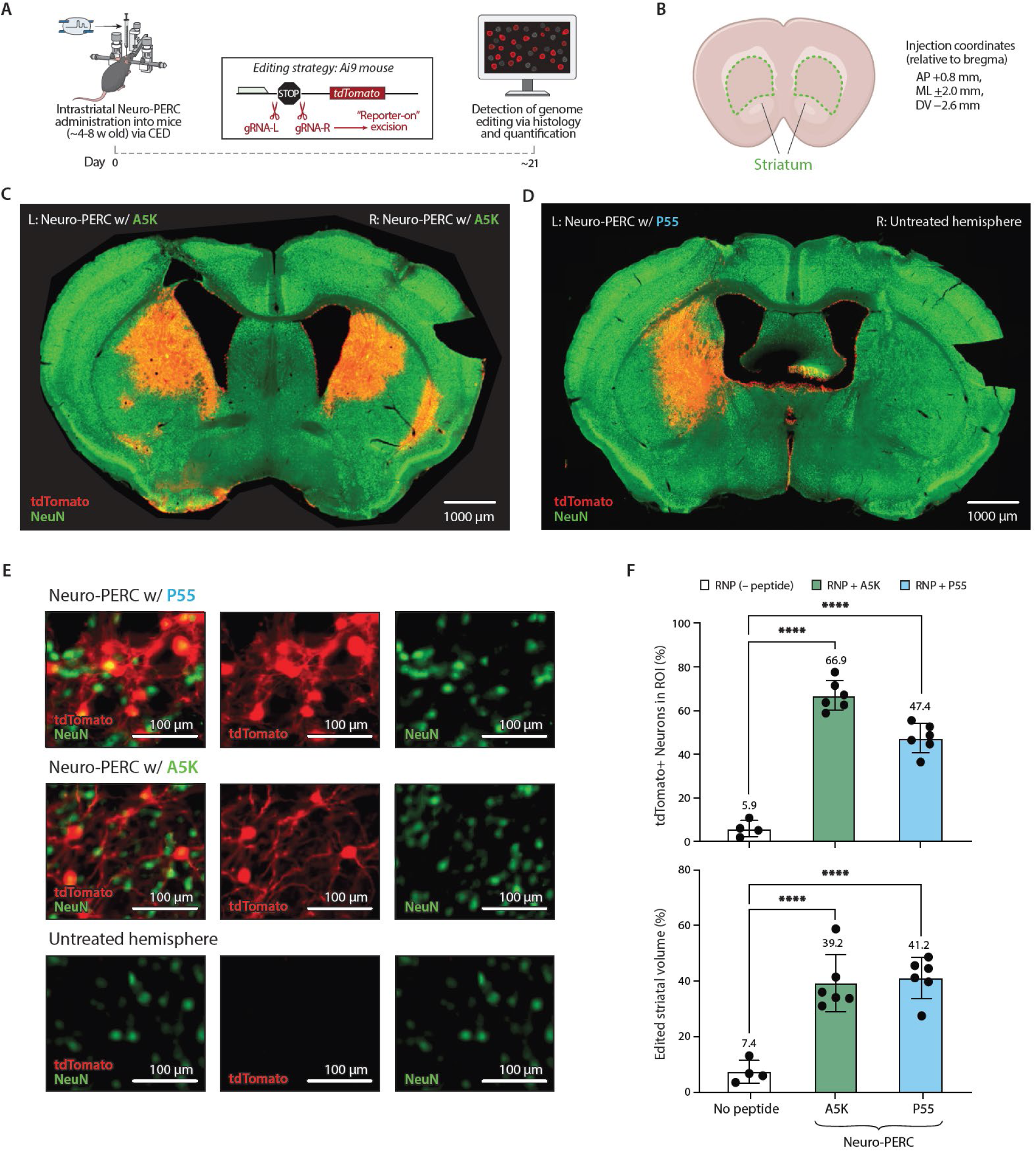
*In vivo* neuronal genome editing in the Ai9 mouse striatum following CED administration of Neuro-PERC. **(A)** Schematic of the *in vivo* mouse experiments. Ai9 mice (∼4–8 weeks old) received intrastriatal administration (either unilateral or bilateral) of Neuro-PERC via convection-enhanced delivery (CED). Genome editing was assessed 3 weeks (unless other timepoint was indicated) post-injection by tdTomato^+^ expression (via histology) resulting from excision of the STOP cassette. **(B)** Coordinates for stereotaxic targeting of the striatum: AP +0.8 mm (anterior to bregma), ML ±2.0 mm (lateral from midline), DV −2.6 mm (depth from skull surface). **(C)–(D)** Representative immunofluorescence microscopy showing tdTomato (red) or GFP (green) expression colocalized with NeuN (neuronal marker; green) in striatal sections from mice treated with A5K- or P55-based Neuro-PERC or untreated hemispheres as controls. Scale bars, 1, 000 μm. **(E)** Representative high-magnification immunofluorescence images of tdTomato^+^/NeuN^+^ neurons in the edited striatal region of Ai9 reporter mice; Single-cell resolution reveals neuronal morphology and local editing patterns within the injection-proximal area. Scale bars, 100 μm. **(F)** Quantification of tdTomato⁺/NeuN⁺ cells within defined regions of interest (ROIs) and measurement of edited striatal volume in Ai9 mice. Each data point represents an individual mouse (n = 4–6 per group). Data are presented as mean ± SD. ****P < 0.0001.

Following CED-mediated intrastriatal administration of Neuro-PERC with peptide A5K, tdTomato^+^/NeuN^+^ neurons occupied 39.4 % ± 5.1% of the striatal volume, with an editing efficiency of 66.9% ± 3.4% within the successfully targeted striatum (P < 0.0001 vs. control for A5K) (Fig. 2C, Fig. 2F, Fig. S2). In an analogous administration of Neuro-PERC with peptide P55, tdTomato^+^/NeuN^+^ neurons occupied 41.2% ± 3.7% of the striatal volume, with an editing efficiency of 49.4 % ± 3.7% within the successfully targeted striatal volume (P < 0.0001 vs. control for P55) (Fig. 2D, Fig. 2F, Fig. S2). Together, these results show that Neuro-PERC mediated robust genome editing across large portions of the striatum, as evidenced by tdTomato⁺ expression. Upon analysis of the region edited by Neuro-PERC (incorporating either peptide) via microscopy, tdTomato^+^ expression localized predominantly to cells with neuronal morphology characteristic of striatal projection neurons and cells that were NeuN^+^ (Fig. 2E). Both formulations achieved comparable striatal coverage, with A5K exhibiting a slightly higher editing efficiency (Fig. 2F). Striatal editing was diffuse following Neuro-PERC administration (with either peptide), as evidenced by tdTomato^+^ expression in adjacent sections and extended beyond the striatum into the globus pallidus (Fig. S3). To assess robustness and reproducibility of the platform, we performed an independent validation of the formulation at the Jackson Laboratory (JAX) using P55-based Neuro-PERC (*50*). Prior murine studies involved Neuro-PERC formulations that were shipped from U.C. Berkeley frozen, with the reagents being thawed at The Ohio State University Medical Center before administration; analogous logistics were employed for the JAX study performed at Bar Harbor, Maine. This study recapitulated our initial findings with peptide P55, with 40.2 ± 2.3 % of the striatal volume edited and an editing efficiency of 38.9% ± 1.8% within the ROI (Fig. S4). Control hemispheres (RNP only) revealed low levels of tdTomato^+^ signal, likely resulting from nominal self-delivery capacity of the three-NLS Cas9 RNP (Fig. S4), which might be predicted based on previous findings (*38*).

We next evaluated Neuro-PERC (incorporating either peptide A5K or P55) in a constitutive green fluorescent protein (GFP) mouse model to establish its versatility using a distinct animal model and editing strategy (Fig. 3A). Cas9 nuclease-mediated GFP knockout was used to quantify editing efficiency and uses a single guide targeting to generate a frame-shifting indel. Using peptide A5K, GFP^−^/NeuN^+^ neurons occupied 51.4 % ± 4.8% of the striatal volume (P < 0.001 vs. control for A5K) with an editing efficiency of 65.4% ± 3.3% within the successfully targeted striatal volume (P < 0.0001 vs. control for A5K) (Fig. 3C, Fig. S5, Fig. 3D). With peptide P55, GFP^−^/NeuN^+^ neurons occupied 48.6% ± 6.0% of the striatal volume (P < 0.01 vs. control for P55), with an editing efficiency of 55.9 % ± 9.7% within the successfully targeted striatal volume (P < 0.001 vs. control for P55) (Fig. S5, Fig. 3D). Editing in the striatum in the GFP mice was diffuse for both peptides, as evidenced by loss of GFP expression across consecutive sections of the injected hemisphere (Fig. S6). Differences in efficiency likely reflect distinct genome editing strategies: GFP knockout requires only a single enzyme-gRNA RNP to reach the nucleus, whereas Ai9 cassette excision depends on paired gRNAs, halving the effective dose of each RNP and necessitating a coordinated pair of double-strand breaks. Despite locus and strategy-dependent variability, these results demonstrate that Neuro-PERC is compatible with multiple murine models and editing approaches.

**Fig. 3:**
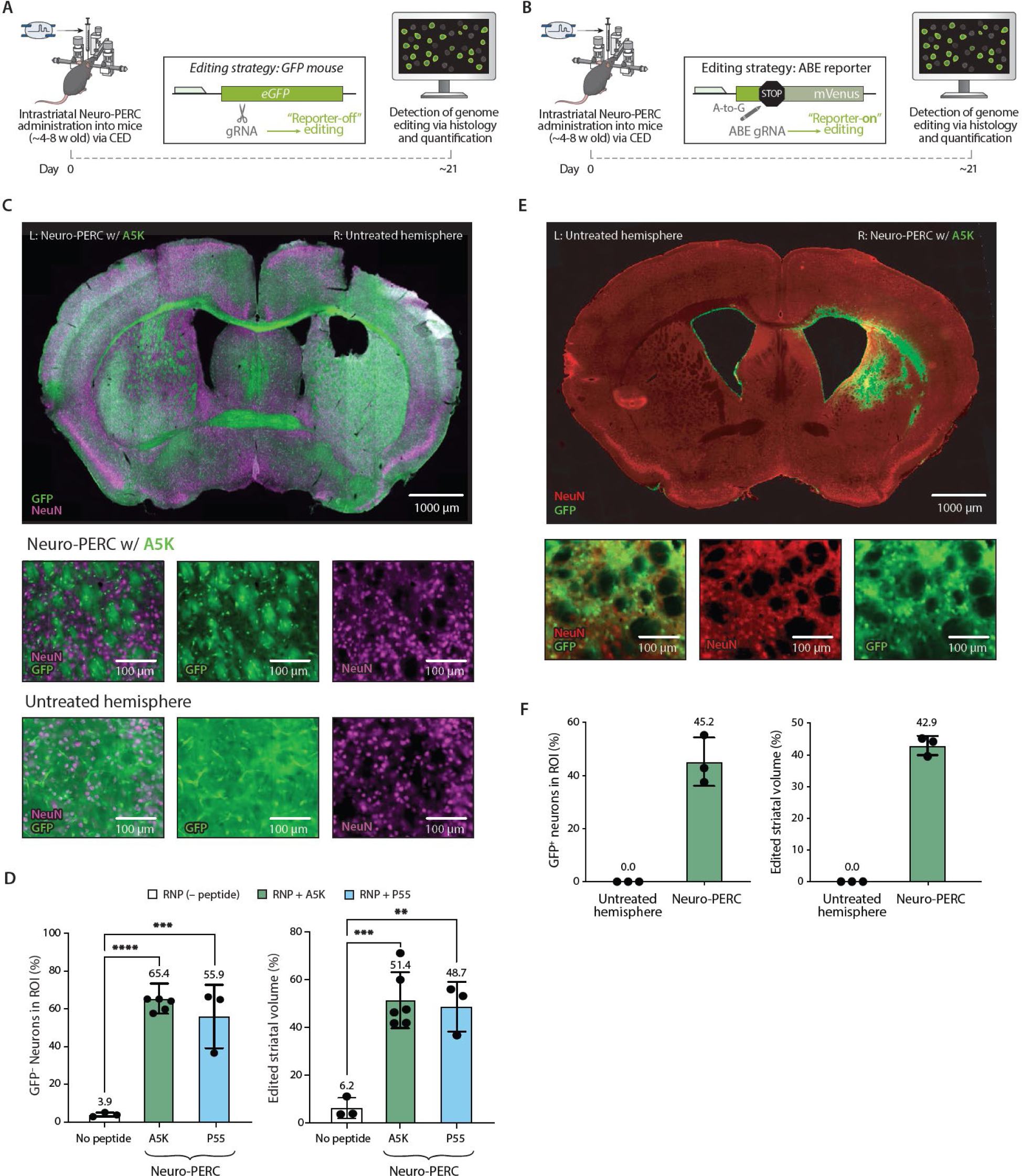
*In vivo* neuronal genome editing in the mouse striatum of other mouse models following CED administration of Neuro-PERC. **(A)** Schematic of the *in vivo* mouse experiments with GFP mice. Transgenic GFP reporter mice (∼4–8 weeks old) received intrastriatal administration (either unilateral or bilateral) of Neuro-PERC via CED. GFP expression in striatal tissue was assessed 3 weeks post-injection (unless otherwise indicated) by the GFP^−^ expression in NeuN^+^ cells. **(B)** Schematic of the *in vivo* mouse experiments with GER10 mice. GER10 mice (∼4–8 weeks old) received intrastriatal administration (unilateral) of Neuro-PERC via CED. GFP⁺ expression in striatal tissue was assessed 3 weeks post-injection (unless otherwise indicated) by quantifying GFP⁺ signal in NeuN⁺ cells. **(C)** Representative immunofluorescence microscopy showing NeuN (magenta) or GFP (green) expression in striatal sections from GFP mice treated with A5K-based Neuro-PERC or untreated hemispheres as controls. Scale bars, 1000 μm. The insets show high magnification representative microscopy of the edited region. Scale bars, 100 μm. **(D)** Quantification of GFP⁻/NeuN⁺ cells within defined ROIs and measurement of unedited striatal volume in GFP mice. Each data point represents an individual mouse (n = 3–6 per group). Data are presented as mean ± SD. **P < 0.01; ***P < 0.001; ****P < 0.0001 **(E)** Representative immunofluorescence microscopy showing NeuN (red) or GFP (green) expression in striatal sections from Ger10 mice treated with A5K-based Neuro-PERC or untreated hemispheres as controls. The insets show high magnification representative microscopy of the edited region. Scale bars, 100 μm. **(F)** Quantification of GFP^+^/NeuN⁺ cells within defined ROIs and measurement of unedited striatal volume in Ger10 mice. Data are presented as mean ± SD. Each data point represents an individual mouse (n = 3 per group).

CRISPR-based adenine base editors (ABEs) (*51*) represent potent members of the genome engineering toolkit, and have been used in clinical trials (*52, 53*). To assess Neuro-PERC’s compatibility with genome editors beyond Cas9, we paired peptide A5K with RNP-format ABE-8e, a potent engineered adenine base editor (*54*). The ABE protein component was further engineered (beyond previously reported embodiments) to increase NLS density, a property that can promote self-delivery of RNP (*38*) or function synergistically with peptide-mediated delivery (*27–29*). We evaluated ABE-8e Neuro-PERC in the GER10 mouse model (*55*), which bears a Venus GFP cassette containing a premature stop codon that can be eliminated via adenine base editing, resulting in a “GFP-on” readout (Fig. 3B). ABE-8e/A5K Neuro-PERC achieved 45.2 ± 1.3% GFP-on across 42.3 ± 1.7% of striatal volume, with diffuse distribution similar to nuclease-based approaches (Fig. 3E, Fig. 3F, Fig. S7), demonstrating that Neuro-PERC mediates brain genome editing with either Cas9 nuclease or ABE across three murine fluorescent reporter models without performing enzyme-specific optimization.

Previous work has shown that Cas9 RNPs elicit a limited immune response following intraparenchymal administration via CED (*17*). With this context, we sought to determine whether Neuro-PERC would provoke an immune response in the brain, assessing this response at different timepoints (21 days and 3 months post-infusion). For both timepoints, the H&E, NeuN, and/or DAPI staining revealed negligible cell loss in the injected hemispheres compared to untreated hemispheres (Fig. S8, Fig. S9, Fig. S10). At both timepoints, we observed a modest immune response in the striatum compared to untreated hemispheres as indicated by Iba1^+^ and GFAP^+^ signal (Fig. S8, Fig. S9). To further investigate, we compared Neuro-PERC formulations bearing either the Ai9-targeting gRNA or a gRNA targeting the murine *Rosa26* locus (henceforth mROSA26), which is not anticipated to have any physiological impact. Notably, we observed comparable GFAP^+^ expression in both hemispheres (Fig. S10), suggesting that the immune response is likely driven primarily by the surgical procedure and/or the presence of Cas9 RNP, rather than by genome editing or tdTomato expression itself. This scenario would be consistent with a prior study wherein Cas9 RNP was likely the principal driver of these immune responses (*17*).

We sought to determine whether the observed inflammation was model-specific by assessing a GFP mouse model at the previously indicated timepoints. In both cases, NeuN and/or DAPI staining revealed negligible cell loss in injected hemispheres compared to untreated controls, with cells appearing uniform and with normal morphology, similar to Ai9 mice (Fig. S11, Fig. S12). At the 3-week timepoint, glial GFAP^+^, and microglial and macrophagic CD68^+^ expression in GFP mice was largely confined to the needle track and the infusion site in the striatum, suggesting the response was primarily driven by mechanical injury from surgery. Notably, at the 3-month timepoint, Iba1^+^ and GFAP^+^ expression was notably reduced compared to the 3-week timepoint and remained localized to the needle entry site in the striatum (Fig. S12), and notably, not expressed in the entirety of edited striatal region. These results establish Neuro-PERC as a well-tolerated CRISPR delivery strategy and a promising candidate for *in vivo* evaluation in larger animal models.

### Neuro-PERC enables efficient, well-tolerated neuronal genome editing in the pig brain

Large animal models are critical for assessing genome editing in anatomically and physiologically relevant systems, helping to de-risk translational efforts. MRI-guided CED allows precise, safe brain infusions with real-time monitoring (*56*) and is used in preclinical and clinical trials of genetic therapies for neurological diseases (*57–60*). We reasoned that combining Neuro-PERC with MRI-guided CED would embody a translationally-relevant delivery strategy for genome editing in a larger animal brain such as a transgenic GFP pig, that constitutively expresses GFP (*61*). Thus, we administered Cas9/A5K-based Neuro-PERC using MRI-guided CED targeting the putamen, a region of the dorsal striatum with anatomical relevance to the human brain (Fig. 4A). The study included one adult pig monitored for three months post-infusion, and a subsequent study with a separate cohort of four 4-week-old pigs evaluated one-month post-infusion.

**Fig. 4:**
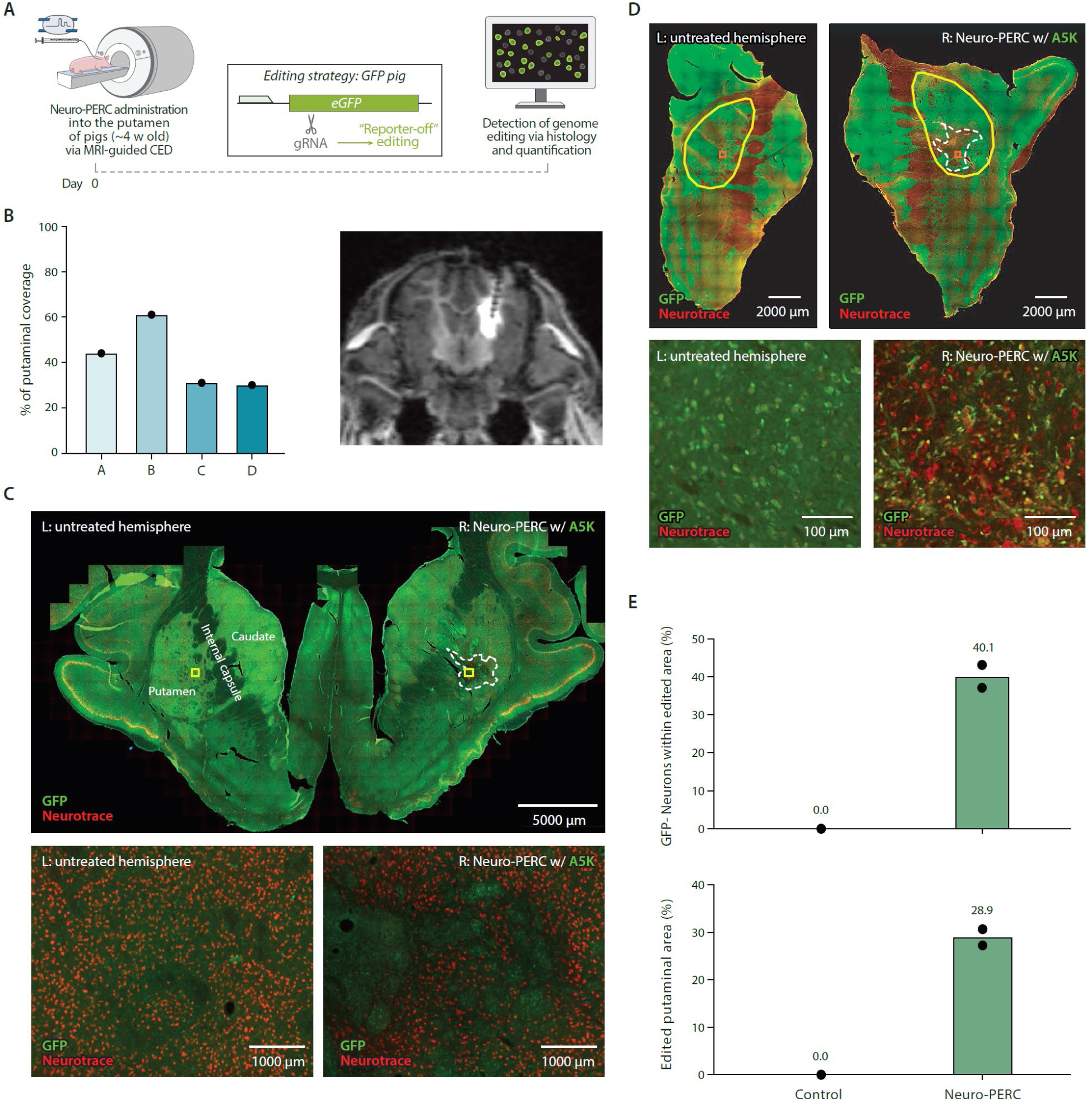
*In vivo* neuronal genome editing in the pig putamen by MRI-guided CED administration of Neuro-PERC. **(A)** Schematic of the experimental procedure showing MRI-guided CED of A5K-based Neuro-PERC into the putamen of GFP-expressing pigs (∼4 weeks old). The editing strategy targeted the eGFP gene using a single guide RNA (sgRNA) to generate a “reporter-off” edit, subsequently assessed by histological and quantitative analyses. **(B)** Percentage of ProHance-infused with Neuro-PERC, localized within the putamen across four 1 month old pigs (A–D). The coronal MRI image (right) illustrates on-target distribution (bright white signal) in pig B, which achieved ∼60% putaminal coverage. **(C)** Brain sections immunostained for GFP (green), NeuroTrace (red), and DAPI (blue), comparing untreated (left) and Neuro-PERC–treated (right) hemispheres. The white dotted outline marks the putamen region showing reduced GFP expression, indicating “reporter-off” editing. Yellow boxes denote areas shown at higher magnification below. Scale bars, 5000 µm (top) and 100 µm (bottom). **(D)** Dissected striatum from untreated and Neuro-PERC–treated hemispheres labeled for GFP (green) and NeuroTrace (red). Yellow outlines indicate the putamen; white dotted lines mark regions with reduced GFP, red boxes denote areas shown at higher magnification below. Scale bars, 2000 µm; insets, 100 µm. **(E)** Quantification of GFP⁻/NeuN⁺ neurons and edited putaminal area one-month post-infusion. Bars represent mean values from two representative adjacent sections of the same animal.

To ensure compatibility with the MRI-guided procedure, the Neuro-PERC formulation was spiked with 2 mM of a gadolinium-based contrast agent (ProHance gadoteridol injection solution) (*62*) to allow for the real-time visualization of the infusion. DLS analysis of Neuro-PERC after freeze-thaw and ProHance addition revealed consistent particle size (∼17 nm), confirming that ProHance is compatible with the platform (Fig. S13). Additionally, a time-course DLS experiment measuring Neuro-PERC size after 8 hours at room temperature confirmed that particle size remained consistent with monodisperse RNP particles (Fig. S1), demonstrating formulation stability required for MRI-guided surgical procedures. Representative samples of Neuro-PERC formulation were collected from the tip of the CED cannula at the start, midpoint, and end of surgery, and SDS-PAGE confirmed formulation integrity throughout the stages of the administration (Fig. S14).

To simulate the surgical context of MRI-guided CED (in pigs or in the clinic), the formulation was maintained in > 3 m of surgical tubing at room temperature for ∼6 hours. Because priming the tubing to remove air is a critical clinical step, we developed priming solution – containing low concentrations of peptide and P-80 – intended to minimize loss of peptide from the Neuro-PERC sample that would chase the priming solution. Because the PERC peptides are amphiphilic, we anticipated their propensity for adsorption to the surgical tubing, and the priming solution was intended to allow an initial bolus of peptide to coat the tubing before the Neuro-PERC formulation entered the tubing. SDS-PAGE analysis of the Neuro-PERC formulation solution remaining in the tubing after priming revealed minimal peptide loss, confirming that the priming procedure did not substantially alter the formulation (Fig. S15).

Upon infusion, ProHance distribution colocalized with 30–61% of the putamen (0.06–0.085 cm³ overlap; putamen volumes 0.139–0.226 cm³) (Fig. 4B, Fig. S13). After performing analysis on representative sections from the pig with the most successful infusion of Neuro-PERC at the one-month timepoint, widespread GFP expression loss was observed throughout the A5K-based Neuro-PERC–treated hemisphere compared to the untreated side (Fig. 4C, Fig. 4D), consistent with efficient editing within the putamen. NeuroTrace^+^ expression revealed minimal neuronal loss or damage within the edited area, with cells appearing morphologically normal in the surrounding tissue as is evident in higher magnification images (Fig. S16). We observed 40.1% of neurons that were GFP⁻/NeuroTrace⁺ (indicating successful editing), representing editing across 28.9% of the putaminal area (Fig. 4E). These findings generally recapitulate our observations in mice and provide strong evidence for cross-species compatibility of the Neuro-PERC platform.

Immunostaining of representative brain sections from the 4-month pig indicated overall normal tissue architecture across both the control and the treated hemisphere, indicative from the DAPI^+^ and NeuN^+^ edited putaminal regions. GFAP^+^ and Iba1^+^ expression revealed localized increases along the infusion tract in the treated hemisphere, similar to what was observed in needle tract from the small animal studies (Fig. S17). Inflammation was not widespread and was limited to the immediate infusion tract, consistent with a localized surgical response. DAPI^+^ and NeuN^+^ staining was consistent throughout the treated hemisphere at both timepoints.

All five pigs exhibited normal behavior throughout the study. Endpoint blood analysis (CBC and CMP) showed no unexpected abnormalities, with minor deviations from baseline consistent with normal surgical recovery, hydration, or transient physiological changes (Table S1). Compared to age-matched naïve pigs, neutrophil, monocyte, and lymphocyte counts remained within normal ranges, indicating no acute or long-term systemic immune response. Slightly elevated platelets on days 4 and 7 likely reflect normal post-surgical variability. Most metabolic panel values were near expected ranges, with minor deviations possibly due to diet or activity, and modest elevation of creatine kinases was not concerning. Genomic DNA from multiple non-target tissues was analyzed by PCR and targeted NGS, revealing no detectable editing and indicating that Neuro-PERC-mediated genome editing was confined to the intended target region (Table S2). Overall, Neuro-PERC was well tolerated with no clinically significant adverse effects.

### Neuro-PERC extends life in an R6/2 murine model of HD

Previous studies have shown that achieving a relatively small proportion (20–30%) of genetically corrected cells can lead to significant improvement in the phenotypic symptoms of Huntington’s disease (HD) (*63, 64*). Building on this evidence and prior success with CRISPR-Cas9 targeting mutant huntingtin protein (encoded by the human m*HTT* gene) in mice (*65, 66*), we applied Neuro-PERC in the well-characterized R6/2 HD model (*67*) that harbors a pathogenic fragment of human m*HTT* and develops early motor deficits at ∼3 weeks of age as well as huntingtin inclusions. Having established the efficiency and tolerability of Neuro-PERC in reporter models, we employed a disease-relevant context, using the R6/2 mouse model of HD to evaluate the therapeutic potential of Neuro-PERC. R6/2 mice (3–4 weeks old, ∼160 CAG repeats) received bilateral striatal infusions of A5K-based Neuro-PERC bearing a gRNA targeting m*HTT* (or a negative control gRNA targeting the mROSA26 locus), after which cohorts were assessed for survival, motor coordination via rotarod and open field tests, and molecular outcomes from striatal microdissection (Fig. 5A).

**Fig. 5:**
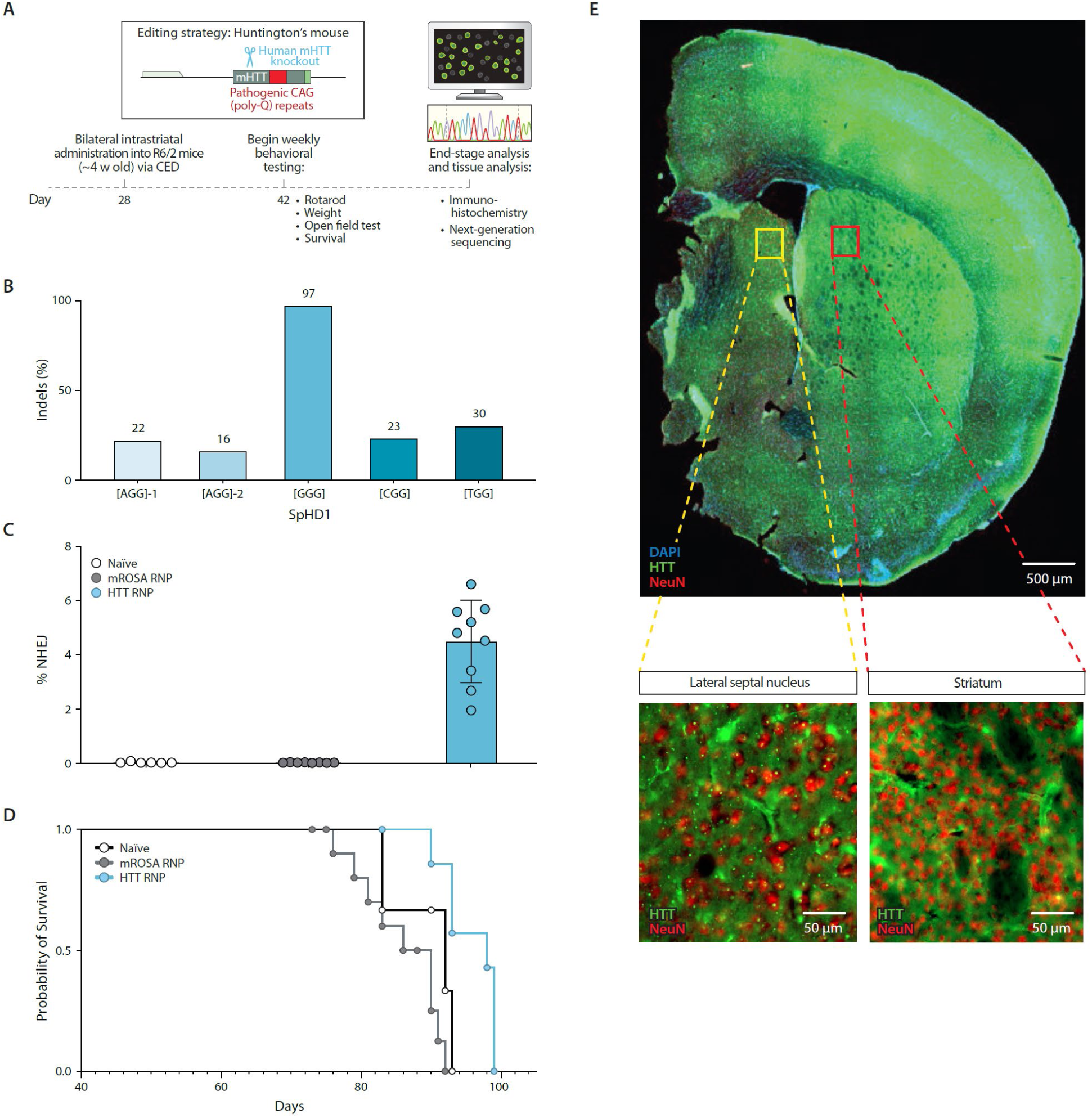
Neuro-PERC can extend life in an aggressive murine model of HD. **(A)** Experimental timeline showing bilateral intrastriatal administration of A5K-based Neuro-PERC delivering Cas9–sgRNA targeting the human mutant huntingtin (mHTT) allele in ∼4-week-old R6/2 mice via CED, followed by behavioral assessments, endpoint tissue analyses and genomic outcomes. **(B)** sgRNA selection in R6/2 fibroblasts by electroporation. The SpHD1 guide ([GGG], shown on the x-axis) achieved the highest editing efficiency, producing 97% indels as determined by Sanger sequencing. **(C)** Quantification of indel frequencies at the mHTT target site in the striatum of treated R6/2 mice. Mice received Neuro-PERC carrying SpHD1 (“HTT RNP,” blue), Neuro-PERC carrying mROSA26 (“mROSA RNP,” gray), or were left untreated (“naïve,” white). Indel frequencies were determined by next-generation sequencing (NGS). **(D)** Kaplan–Meier survival curves for R6/2 mice treated as in (C). Statistical analysis by Log-rank (p = 0.0003) and Gehan–Breslow–Wilcoxon (p = 0.0027) tests revealed significantly extended lifespan in HTT-targeted Neuro-PERC–treated mice versus controls. **(E)** Representative striatal section from an NeuroPERC-treated mouse stained for mutant huntingtin (mHTT, green), NeuN (red), and DAPI (blue). The lateral septum nucleus served as an internal control. Reduced mHTT colocalization with NeuN (yellow) indicates decreased protein expression. Scale bars, 500 μm; 50 μm (insets).

To adapt an established *S. aureus* Cas9 strategy for human huntingtin knockout (*65*), *S. pyogenes* Cas9-compatible gRNAs were screened via RNP electroporation into R6/2-derived fibroblasts, identifying SpHD1, featuring a GGG protospacer adjacent motif (PAM) – which induced insertions and deletions (indels) with an efficiency of 97% – as a highly potent guide for efficient m*HTT* knockout by targeting human *HTT* exon 1 (Fig. 5B, Table S3). Based on its robust editing efficiency and knockout of m*HTT* in R6/2-derived fibroblasts, SpHD1 RNP was selected to advance to *in vivo* Neuro-PERC studies (henceforth described as HTT RNP).

Genomic DNA was isolated from RNP-infused striatal tissue for deep sequencing of the m*HTT* gene, revealing ∼4% indels in HTT RNP-treated mice versus controls (Fig. 5C). This outcome generally aligns with our expectations from the murine studies discussed above and is consistent with those results via the following calculation, which incorporates estimates of typical coverage and efficiency observed in reporter mice: 40% area/coverage × 40% editing efficiency × 25% of cells eligible for editing (based on the 3:1 glia-to-neuron ratio estimated in the striatum (*68*)), yielding an anticipated editing efficiency of approximately 4%. Notably, this indel frequency is comparable to that reported in a prior study where AAV-mediated *S. aureus* Cas9 delivery partially rescued R6/2 mice and extended their lifespan (*65*).

Survival analysis revealed that m*HTT* knockout extended the lifespan of treated mice compared to both naïve and control group which received the mROSA26 gRNA (Fig. 5D). The treated cohort survived ∼9 days longer than the control gRNA cohort, indicating a shift toward extended survival in the m*HTT*-targeted gRNA cohort (Log-rank, *p* = 0.0003; Gehan–Breslow–Wilcoxon, *p* = 0.0027). The lifespan of mice treated with m*HTT*-targeted Neuro-PERC ranged from 83−99 days, compared to 73−90 days for control mice and 83−91 days for naïve animals. Given the aggressive disease progression in the R6/2 model, this lifespan extension represents a meaningful benefit, highlighting the potential of our Neuro-PERC intervention.

To assess the functional outcomes of our intervention, we conducted the rotarod test and the open field test and monitored body weight of the mice. Rotarod performance showed no improvement with m*HTT*-targeting Neuro-PERC compared to either naïve or control-gRNA cohorts at either timepoint tested (Fig. S18). Body weight decreased similarly across groups over the course of the study, suggesting that the intervention did not negatively impact the weight of the mice as compared to the controls (Fig. S18). Overall, no overt behavioral deficits were ameliorated by our intervention within the timeframe of this study but – importantly – the treatment did not worsen outcomes compared to naïve or control cohorts.

Immunohistochemistry was performed on brain sections from treated and control mice to assess mutant huntingtin protein (mHTT) expression alongside NeuN and DAPI staining (Fig. 5E, Fig. S19, Fig. S20). The lateral septum nucleus served as an internal reference region within each hemisphere. Treated mice exhibited a marked reduction of mHTT signal in the striatum relative to the lateral septum, whereas control mice displayed uniform mHTT expression across regions, confirming successful region-specific knockout of mHTT in the treatment group.

The results from this experiment demonstrate that Neuro-PERC can be utilized to suppress expression of the m*HTT* gene in the R6/2 mouse following intrastriatal administration via CED, extending lifespan in a severe model of Huntington’s disease.

## DISCUSSION

*In vivo* administration of CRISPR-Cas9 for genome editing of the brain holds tremendous promise for advancing much-needed therapeutic strategies across various neurological diseases and can potentially mitigate neurodegeneration (*69, 70*). However, safe and effective delivery of CRISPR effectors to the brain remains a major challenge in the field of therapeutic genome editing and is the single greatest impediment to clinical translation (*71*). In this study, we adapted the non-viral PERC technology (*5*) to establish Neuro-PERC, an optimized delivery platform that mediates efficient *in vivo* neuronal editing in both small and large animal models. Neuro-PERC facilitates efficient genome editing of the brain while minimizing immune response in reporter animal models. This approach addresses several key challenges associated with *in vivo* genome editing in the brain, exemplified by its capacity to prolong survival in a severe mouse model of the neurodegenerative condition Huntington’s disease as well as its successful deployment in a large animal via MRI-guided CED – as is routinely performed for clinical administration of genetic therapies for the brain. These findings establish *in vivo*-optimized peptide-mediated delivery of CRISPR enzymes in RNP format – Neuro-PERC – as a promising candidate for clinical genome editing in the brain.

The main challenge in adapting PERC for *in vivo* brain editing was developing formulations that yield monodisperse nanoscale particles with amphiphilic peptides for efficient intracellular delivery, while also demonstrating effective genome editing in neural cells. The resulting Neuro-PERC formulations are highly robust, maintaining consistent small particle size after freeze–thaw cycles and remaining fully compatible with sterile filtration. In NPCs, Neuro-PERC mediated highly efficient genome editing rates that far surpassed those attained using RNP-only controls, confirming its capacity to edit neuronal cells. Beyond the potent editing performance, the PERC platform offers significant translational advantages: it is simple and straightforward to manufacture, requiring only the mixing of three well-established macromolecular components, followed by filtration. This streamlined process promotes consistent quality, scalability, and reproducibility. Importantly, the accessibility of cGMP-grade RNP – already used in multiple clinical trials and in the first approved CRISPR-based therapeutic – provides a strong foundation for translation of Neuro-PERC for clinical applications.

A major consideration in deploying Neuro-PERC *in vivo* was ensuring compatibility with clinically relevant administration techniques. The small, monodisperse Neuro-PERC particles are ideally suited for direct intraparenchymal delivery using convection-enhanced delivery (CED), enabling both robust local effector concentrations and broad distribution within the striatum. In murine models, Neuro-PERC administered by CED achieved efficient and reproducible neuronal editing. Independent validation performed by a distinct laboratory confirmed these results, underscoring reproducibility in multiple research settings. Neuro-PERC also supports diverse genome editing strategies, including generation of indels to induce frameshift and protein knockout, programmed excision, and adenine base editing, all without the need for enzyme-specific optimization. The ability of Neuro-PERC to reproducibly deliver distinct CRISPR effectors with high efficiency and broad tissue coverage across multiple preclinical models highlights its strong translational potential.

Neuro-PERC exhibited a tolerable immune profile following intraparenchymal administration via CED at multiple timepoints. Across multiple mouse models, including Ai9 and GFP reporter strains, NeuN^+^ and DAPI staining revealed negligible neuronal or cell loss, with cells appearing morphologically normal compared to untreated controls. Modest immune activation, as assessed by Iba1^+^, GFAP^+^, and CD68^+^ staining, was largely confined to the needle track and diminished over time, with lower prevalence at three months compared to three weeks post-injection. The immune response to Neuro-PERC is comparable to the previously reported immune response to a peptide-free administration of Cas9 RNP, suggesting that the PERC peptides do not make a substantial contribution to the immune response (*48*). These results indicate that Neuro-PERC elicits only localized and transient immune responses, which bodes well for tolerability in potential therapeutic applications in the brain.

Evaluation of candidate genetic therapies in large mammals such as pigs provides valuable insights into the scalability of candidate and the therapeutic potential of the strategies for translation for neurological diseases for the human brain (*67*), and we have demonstrated the first precision genome editing in a large animal brain mediated by a non-viral delivery platform. Pigs offer distinct utility regarding assessing scalability and delivery as their striatal structure is comparable to that of humans. Pigs thus provide a preclinical model that bridges the anatomical and physiological gap from rodent models to clinical use (*74*). In addition, pig studies provide insight into the cross-species compatibility of Neuro-PERC. Using MRI-guided CED (*75, 76*) to deliver Neuro-PERC into the putamen of pigs that constitutively express GFP, we achieved efficient neuronal editing. Importantly, Neuro-PERC remained stable throughout the multi-hour procedure, including extended exposure to surgical tubing and contrast agent, demonstrating robustness in a clinic-like context. The surgical procedure closely resembled human neurosurgical protocols, further supporting translational relevance. This study also established compatibility of Neuro-PERC with both murine and porcine neurons and demonstrated that a ∼15× scale-up of reagents was feasible, reinforcing its potential for clinical translation. Neuro-PERC was also successful in editing human iPSC-derived neurons, suggesting broad compatibility with mammalian neurons.

Huntington’s disease results from pathogenic expansion of trinucleotide repeats in the huntingtin gene (*HTT*), causing progressive motor and cognitive decline due to degeneration of striatal medium spiny neurons. Neuro-PERC was used to rescue a murine model of HD. Treatment extended lifespan, demonstrating that mHTT suppression provides therapeutic benefit. Immunohistochemistry confirmed reduced mHTT without neuronal loss, and behavioral and weight analyses indicated good tolerability. These results highlight Neuro-PERC’s potential as a non-viral, targeted platform for suppressing disease-causing genes *in vivo*, providing both survival benefit and partial rescue of molecular pathology in neurodegenerative disease model. This study also highlights the platform’s potential to enable a non-viral, targeted, and disease-modifying intervention for neurodegenerative diseases such as HD.

In summary, Neuro-PERC represents a robust and clinically relevant platform for non-viral CRISPR delivery to the brain, with potential to address the longstanding challenge of safe and efficient *in vivo* genome editing of neural tissue. Its capacity to generate monodisperse, stable particles that support efficient neuronal editing across both murine and large animal models – while exhibiting a tolerable host response – marks a key advance for therapeutic genome editing. Compatible with clinically relevant CED administration and multiple classes of CRISPR enzyme, Neuro-PERC is a versatile, scalable, and well-tolerated platform. These attributes position Neuro-PERC as a promising candidate for clinical translation in neurotherapeutics.

## MATERIALS AND METHODS

### Study design

Neuro-PERC containing 50 µM Cas9 RNP paired with 200 µM delivery peptide (total volume 7 µL per hemisphere; 350 pmol RNP) either A5K or P55 as administered into the striatum of reporter mice via CED to assess *in vivo* genome editing efficiency. The formulation was subsequently scaled for large-animal studies. In a pilot *in vivo* pilot large animal experiment, a single adult pig (NSRRC:0016 GFP NT92) received a unilateral infusion of Neuro-PERC containing 54 µM RNP and 200 µM A5K peptide (total volume 60 µL; 3, 240 pmol RNP) into the right putamen using real-time magnetic resonance imaging (MRI)-guided CED. The animal was monitored longitudinally for three months to evaluate systemic and long-term safety. In a subsequent study, a cohort of four 4-week-old pigs (NSRRC:0016 GFP NT92) received unilateral infusions of A5K-based Neuro-PERC into the putamen (108–110 µL per hemisphere) via MRI-guided CED and were humanely euthanized approximately one-month post-infusion. Blood samples were collected before surgery and at designated timepoints throughout the experimental period to assess systemic and immune responses.

### Cas9 proteins

The *S. pyogenes* Cas9-triNLS cysteine-free variant (C80S, C574S; Addgene #196244) was expressed and purified by the Wilson laboratory following established protocols. Purified Cas9 was concentrated to ∼50 µM in storage buffer (25 mM sodium phosphate pH 7.25; 150 mM NaCl pH 7.5, 10% glycerol) and stored at −80°C.

### Purification of Spy-tagged TadA-8e and SpyCatcher-nCas9 proteins

Plasmids encoding C-terminal SpyTag–TadA-8e or N-terminal SpyCatcher–nCas9 were transformed into BL21 STAR DE3 or Rosetta DE3 cells, respectively. 4–5 colonies were grown overnight in 100 mL Luria-Bertani (LB) medium with the appropriate antibiotic. For expression, 30 mL of overnight culture was added to 1 L terrific broth (with 100 µM ZnCl₂ for TadA-8e), grown to OD ∼2 (TadA-8e) or ∼1.5 (nCas9), and induced with 0.5 mM or 0.2 mM IPTG at 16°C for 24 h or 16–18 h, respectively. Cells were harvested, resuspended in the respective lysis buffer (20 mM HEPES pH 7.5, 10% glycerol, 1 mM TCEP; 300 mM NaCl, 20 mM imidazole for TadA-8e; 1 M NaCl, 20 mM imidazole, 0.1% Triton for nCas9) with protease inhibitors, and sonicated. Lysates were clarified and loaded onto tandem HisTrap FF columns (Cytiva), washed, and eluted with 300 mM imidazole. Eluates were further purified on tandem heparin columns using a 0.3–1 M NaCl gradient on an ÄKTA Pure system. Proteins were concentrated (10 kDa MWCO for TadA-8e, 100 kDa MWCO for nCas9), flash-frozen in liquid nitrogen, and stored at −80°C.

### Conjugation of TadA-8e with nCas9

Purified TadA8e was conjugated with purified nCas9 at a 4× excess molar ratio for 30 minutes at room temperature on a shaking platform. The mixture was then concentrated to a volume of ∼ 3 mL using a 100 kDa MWCO Amicon filter (Millipore Sigma) and loaded over a Hi-Load GF16/60 S200 Superdex column (Cytiva) that had been pre-equilibrated in IEX-A buffer. Purified protein was then concentrated using a 100 kDa MWCO Amicon filter (Millipore Sigma) to 40–50 μM and filtered with 0.22 µm centrifugal filters (Corning, USA)

### RNP formation

The *S. pyogenes* single guide RNAs (sgRNA) were with manufacturer-recommended standard chemical modifications from IDT (Coralville, IA) and resuspended in diethyl pyrocarbonate (DEPC)-treated water. Spacer sequences and PAM for each gRNA used is listed in Table S3. sgRNA stocks (100 µM) were diluted to a final concentration of 37.5 µM in 1× Folding Buffer (10× folding buffer: 200 mM HEPES-NaOH pH 7.5, 1.5 M NaCl). sgRNAs were denatured at 95°C for 5 min and allowed to gradually cool to room temperature (RT) on the bench (∼25 min). The Cas9 protein was diluted to 25 µM in 1× GF buffer supplemented with 200 mM trehalose (20 mM HEPES-NaOH pH 7.5, 150 mM NaCl), and MgCl₂ was added fresh to 2 mM immediately before use, yielding a final concentration of 1 mM upon RNP complex formation. For RNP complex formation, folded sgRNA (37.5 µM) and diluted Cas9 (25 µM) were combined at a 1:1 volume ratio, corresponding to a 1.5:1 molar ratio (sgRNA:Cas9) and yielding a final RNP concentration of 12.5 µM based on Cas9 protein. Cas9 was added slowly into sgRNA with gentle mixing, and was incubated at 37°C for 10 minutes before being maintained at RT for immediate use, placed on ice for short-term handling, or flash-frozen for storage.

### Peptide synthesis

The peptide A5K and P55 (CPC Scientific) was synthesized via custom solid phase synthesis with 95% purity. All peptides were stored lyophilized or as 10 mM stocks in DMSO at −20°C in a desiccator. The peptide sequences of A5K and P55 are indicated in Fig. 1B.

### Preparation of *ex vivo* and Neuro-PERC formulations

For *ex vivo* PERC formulations, peptides (10 mM in 100% DMSO) were diluted in DEPC-treated water to 1 mM (resulting in a solution of 90% water and 10 % DMSO) and added to pre-formed RNP. In all cases of peptide-enabled delivery *in vitro*, the final concentration of DMSO was proportional to the peptide concentration, at 0.1% DMSO per 10 µM peptide. The same concentration of DMSO was used for the corresponding RNP only negative control conditions.

For *ex vivo* formulations including P-80, were prepared by mixing 1 µL of 10 mM peptide in 100% DMSO (resulting in 1 mM peptide) with 1 µL of 5% P-80 stock (P-80 is originally at 10%), 3 µL of water, and 5 µL of 1× GF buffer, yielding 10 µL of peptide-P-80 intermediate. An aliquot of the peptide-P-80 intermediate was mixed with pre-formed RNP in 1× GF buffer and combined by mixing with a pipette, yielding a final formulation that was subsequently added to media. The peptide: RNP molar ratio was maintained at 20:1 in all *in vitro* experiments unless otherwise indicated.

For *Neuro-PERC* formulations, RNP was prepared at 12.5 µM based on Cas9 protein and concentrated using 0.22 µm centrifugal filters (Corning, USA). Filters were first equilibrated with 1× GF buffer by centrifugation at 1, 000×g, and residual buffer was removed by inversion and re-spinning at the same speed. RNP was loaded onto the pre-wet filters and concentrated to the desired volume by centrifugation at 13, 000× g in multiple short intervals, gently resuspending the retentate after each spin to ensure homogeneity. Spin durations were increased as volume decreased, repeating cycles until the desired concentration was achieved. Samples were recovered by inverting the filters into pre-weighed sterile tubes and centrifuging at 1000× g, and final volumes were determined gravimetrically. Protein concentrations were quantified spectrophotometrically at 280 nm. For peptide addition, a peptide-P-80 intermediate was prepared by first diluting P-80 in water, then adding the DMSO-containing peptide. An example of this preparation for the material for the pig experiment for a total volume of 300 µL : Neuro-PERC was prepared at a total volume of 300 µL while maintaining the same composition as the mouse-scale formulation (final concentrations: 200 µM peptide, 2% DMSO, 0.06% P-80; 4:1 peptide: RNP molar ratio, assuming 50 µM RNP). A 25 µL peptide premix was prepared by combining 6.0 µL of 10 mM peptide (in 100% DMSO), 13.8 µL of 1.25% P-80, and 5.2 µL of water. The premix was diluted 1:1 by volume with 1× GF buffer to generate a 50 µL working cocktail containing 1.2 mM peptide, 12% DMSO, and 0.35% P-80. One part of this cocktail (50 µL) was then combined with five parts concentrated RNP (250 µL at 50 µM) to yield 300 µL of Neuro-PERC formulation with the desired final concentrations. Prior to use, the Neuro-PERC and other indicated samples were sterile filtered through pre-wetted (with 1× GF buffer) 0.22 µm centrifugal filters (Corning, USA). The Neuro-PERC formulation was then maintained at RT for immediate use, placed on ice for short-term handling, or flash-frozen for storage.

### Dynamic light scattering

Representative samples (which contained 17 µL of GF buffer with 3 µL of Neuro-PERC stock solution to a final volume of 20 µL) were assessed using a Zetasizer Nano ZS (Malvern Panalytical) instrument with a quartz low-volume cuvette (ZEN2112). The sample was incubated at RT for 1 minute and mixed by pipetting. Control samples were subjected to identical handling and volume conditions. Particle size was measured at room temperature, and data were analyzed ‘by number’ using Zetasizer analysis software. Each sample was analyzed three times, resulting in a histogram generated from 14 individual measurements. All size distribution values are reported in nanometers (nm).

### Transmission electron microscopy (TEM)

Samples were prepared by pipetting 10 μL of 0.1 µM RNP solution onto a copper grid (Electron Microscopy Services, CF400-CU). Grids were allowed to incubate at room temperature for two minutes, then washed with DEPC-treated water. The grids were then exposed to 1% uranyl acetate washed with DEPC-water and blotted dry. Particle size was quantified using ImageJ. To determine the average particle size and size distribution, at least four images per condition were accessed with at least 10 particles per image. The particle size data was curve fit using a gaussian distribution in GraphPad Prism (version 5.02; GraphPad Software, San Diego, CA, USA).

### Cell culture with NPCs from Ai9 mice

NPCs were isolated from the cortical tissue of embryonic day 13.5 Ai9-tdTomato homozygous mouse embryos. Cells were cultured as non-adherent neurospheres in NPC medium consisting of DMEM/F12 with GlutaMAX supplement (Thermo Fisher), B-27 supplement (no. 12587-010), N-2 supplement, MEM non-essential amino acids (no. 11140-050), 10 mM HEPES, 1, 000× 2-mercaptoethanol, and 100× Penicillin-Streptomycin. The medium was further supplemented with 20 ng/mL each of FGF-basic (BioLegend) and EGF (Thermo Fisher). Growth factors were replenished every three days, and cultures were passaged every six days using the Neural Tissue Dissociation Kit (Miltenyi Biotec). NPC identity was confirmed by immunostaining for GFAP and Nestin, and cultures were routinely tested for mycoplasma contamination.

### Direct delivery of Neuro-PERC to NPCs

The day before passage, 96-well plates were coated with a mixture of 100 μg/mL poly-DL-ornithine (Sigma-Aldrich), 5 μg/mL laminin (Sigma-Aldrich), and 10 μg/mL fibronectin (Sigma-Aldrich). On day 6, dissociated cells were plated onto the pre-coated plates with 5, 000 cells per well in 100 µL of media. Media was refreshed on the day of treatment, and cells were processed for analysis via flow cytometry at three days post-treatment.

### Flow cytometry

Flow cytometry was performed on an Attune NxT flow cytometer with a 96-well autosampler (Thermo Fisher). Cells were pelleted (1, 000×g, 3 min), washed with PBS, and resuspended in FACS buffer (PBS with 2% FBS and 1 mM EDTA). Cell viability was assessed using the Fixable Violet Dead Cell Stain Kit (1:1, 000; Thermo Fisher, #L34964) for 20 min at 4°C in the dark. After staining, cells were washed, treated with 0.25% trypsin/EDTA (Thermo Fisher) for 10–15 min, and dissociated with DNase (2 µL/10 mL PBS; Neurosphere Dissociation Kit, Miltenyi Biotec). Trypsin was neutralized with DMEM containing 10% FBS, and single-cell suspensions were analyzed using the Attune NxT VL1 and YL1 channels. Data were processed in FlowJo (BD Biosciences).

### Cell line generation and differentiation

The Kolf2.1J1 stable iPSC cell line (JAX) containing a knock-in of an inducible NGN2 motor neuron transcription factor transgene cassette in the CLYBL safe-harbor locus was generated using SpCas9. Cells were confirmed negative for mycoplasma contamination (quarterly testing, WiCell) and exhibited a normal karyotype. iPSCs were differentiated into motor neurons using a previously described protocol (*45*).

### Direct delivery of Neuro-PERC to iPSC-derived neurons

All experiments were performed using induced motor neurons 10 days of induction with doxycycline. Cells were plated in 96-well plates at a density of approximately 20, 000 cells per well and maintained in day 10 differentiation medium prior to treatment. Neuro-PERC with a gRNA targeting B2M (provided in Table S3) w was added directly to the culture media, with each condition performed in triplicate. Each replicate was processed independently for downstream sequencing analysis. Live-cell imaging was performed using an Incucyte Live-Cell Analysis System (Sartorius) to assess cytotoxicity. Genomic DNA was extracted at defined timepoints using QuickExtract DNA Extraction Solution (Lucigen, no. QE09050) according to the manufacturer’s instructions and stored at −20°C prior to sequencing. PCR amplification (primers provided in Table S6) was performed in 20 µL reactions containing Biomix Red (Bioline, no. BIO-25006) and 1 µL genomic DNA. Thermocycler conditions were as follows: 95°C for 3 min; 40 cycles of 95°C for 30 s, 58°C for 30 s, 72°C for 1 min; followed by a final extension at 72°C for 5 min. PCR products were verified on a 2% agarose gel, and editing efficiency from three biological replicates was quantified by Sanger sequencing (Quintara Biosciences) using the forward primer. Indel rate was quantified using ICE analysis.

### Small animal surgical procedures

All animal procedures complied with the Animal Welfare Act and NIH guidelines and were approved by the IACUC at The Ohio State University (mice), University of Missouri (pigs), or University of California, Berkeley (mice). Ai9 mice (B6.Cg-*Gt(ROSA)26Sortm9(CAG-tdTomato)Hze*/J strain# 007909), constitutive GFP mice (C57BL/6-Tg(CAG-EGFP) 1Osb/J Strain #:003291 Hemizygous), or GER10 (C57BL/6N-*Gt(ROSA)26Sorem3(CAG-Venus*)Rkuhn*/MurrMmjax strain #069724-JAX) mice were group-housed on a 12 h light-dark cycle with ad libitum food and water; room temperature and humidity were 21 ± 2°C and 50–60%, respectively. Mice were anesthetized with 2% isoflurane (Baxter, Deerfield, USA), given subcutaneous meloxicam and buprenorphine (Buprenorphine ER-Lab), and placed in a stereotaxic frame. A burr-hole was drilled at striatal coordinates relative to bregma (AP +0.8 mm; ML ±2.0 mm; DV –2.6 mm). A fused silica cannula (1-mm step) was used for unilateral CED; 7 µL of formulation was infused at 1 µL/min using a microsyringe pump (World Precision Instruments). The syringe remained in place for 2 min before removal. Incisions were closed with sutures or wound clips, and a topical antibiotic applied. Mice were kept at 37°C, Lubrifresh P. M. ointment (Major, Indianapolis, IN, USA, NDC 0904-6488-38) was applied to the eyes, and animals were monitored until recovery, then evaluated twice daily for 5 days.

### Large animal surgical procedures

Animals were placed prone in an MRI-compatible stereotactic frame with flex coils (Flex Small 4, Siemens), and a burr hole craniotomy was performed. A skull-mounted, MRI-compatible ball-joint guide array (BJGA) was positioned approximately 20 mm posterior to the eye and 11 mm lateral to the suture. Animals were moved into a 3T MRI scanner to calculate and optimize the trajectory to the caudate, avoiding critical vasculature and ventricles. A ceramic, custom-designed fused silica reflux-resistant cannula with a stepped tip (Clearpoint) was used for infusion. For real-time visualization, the Neuro-PERC formulation was co-administered with gadoteridol (2 mM, Prohance; Bracco Diagnostics, Monroe Township, NJ). Infusion rates were gradually increased up to a maximum of 3 µL/min. Upon completion, the BJGA was removed, hemostasis was achieved, and a titanium burr hole cover was placed. Animals were returned to home cages and monitored during recovery. Follow-up MRI was performed at 1 week and prior to the experimental endpoint. Routine blood was collected from animals at a volume of 2–5 mL/sample via venipunture of cranial vena cava, jugular vein, or other peripheral vessel at indicated timepoints. Animals were humanely euthanized after 4 weeks or 3 months, and brains were harvested.

### Tissue collection and processing

Animals were humanely euthanized at the study endpoints and transcardially perfused with phosphate buffered saline (PBS, mice) or heparinized saline (pigs), followed by 4% buffered paraformaldehyde (PFA), pH 7.4. Brains were harvested, post-fixed for 24 hours (mice) or 72 hours (pigs), blocked in 6 mm coronal sections (pigs), and stored in 30% sucrose in PBS (w/v) until sectioning. Tissue was then frozen in optimal cutting temperature (−18°C) tissue freezing medium (Leica) and cut into 40 μm serial, coronal sections with a sliding microtome and stored in antifreeze (glycerol/ethylene glycol/PBS) until they were processed for immunohistochemistry.

Brain hemispheres were fixed in 4% paraformaldehyde (PFA; Electron Microscopy Sciences, #15710), cryoprotected in 30% sucrose, embedded in optimal cutting temperature (OCT) compound (Thermo Fisher, #23-730-571), and stored at −80°C. Coronal sections (30 µm or 40 µm) were prepared using a cryostat (Leica CM3050S) and mounted onto positively charged slides (Fisherbrand, #1255015).

For immunostaining, free-floating or mounted sections were washed in PBS containing 0.3–0.5% Triton X-100 (Fisher Scientific) and blocked in either 5% goat serum (MilliporeSigma, G9023) with 1% bovine serum albumin (MilliporeSigma, F0850) or 1× Animal-Free Blocker (Vector Laboratories, SP5030). Sections were incubated overnight at 4°C in the appropriate primary antibodies (provided in Table S4) diluted in blocking buffer, followed by incubation with the corresponding secondary antibodies (provided in Table S5) for 1 h at room temperature. Nuclei were counterstained with DAPI (1:5, 000 in PBS; MilliporeSigma, MBD0015), washed in PBS, and mounted with Fluoromount (MilliporeSigma, F4680). Coverslips were sealed with clear nail polish (Electron Microscopy Sciences, #72180).

Fluorescent imaging was performed using a Zeiss Axioscan 7 (20× objective; Zeiss, Oberkochen, Germany) for automated whole-slide scanning and confocal microscopes including the Evos Revolve (Echo, San Diego, CA), Leica Stellaris 5 (Leica Microsystems, Deer Park, IL), and Zeiss LSM 710 AxioObserver (Zeiss, Oberkochen, Germany), each equipped with 10× or 20× objectives. For each striatum, approximately five images were acquired at 20–25× magnification across multiple sections. Z-stack imaging (4–6 stacks per field, 1024 × 1024 px) was used to quantify protein and neuronal marker expression. Images were processed in ZenLite (Zeiss Zen 3.10), ImageJ, or QuPath with consistent thresholds applied across treatment groups.

For striatal analysis of the pig brain, coronal pig brain slices were prepared from post-fixed brains, and the striatum was carefully dissected from each slice using a sterile scalpel. Tissue was cryoprotected in 30% sucrose in PBS, embedded in OCT compound, and frozen. Coronal sections (40 μm) were cut using a sliding microtome and stored in antifreeze solution until immunostaining. Free-floating sections were washed in PBS containing 0.3% Triton X-100, post-fixed in 10% neutral buffered formalin, and blocked in 5% bovine serum albumin in PBS/0.3% Triton X-100. Sections were incubated with anti-GFP primary antibody (1:200; Invitrogen) for 48 hours at 4°C, followed by Alexa Fluor 546-conjugated secondary antibody (1:300) for 1.5 hours at room temperature. Deep Red NeuroTrace (1:200) was applied for 1 hour at room temperature, and sections were treated with Sudan Black B in 70% ethanol to reduce autofluorescence. Nuclei were counterstained with DAPI or NucBlue, and sections were mounted with ProLong Gold antifade mounting medium.

### Image Analysis

Image analysis for base genome editing and Ai9 genome editing was performed in QuPath (v0.6.0; University of Edinburgh). Regions of interest (ROIs) outlining each striatum were manually drawn using the polygon tool to annotate both the striatal borders and the inner tdTomato^+^ or GFP^+^ edited regions. For base editing, the green channel (GFP) defined edited regions; for Ai9 editing, the red channel (tdTomato, tdT) was used. Cell detection in both analyses used the red (NeuN) channel with optimized neuronal parameters (background radius = 20 px, sigma = 1 px, minimum area = 20 µm², maximum area = 400 µm², cell expansion = 5 px, intensity threshold = 5). Two-channel quantification identified GFP⁺ or tdT⁺ neurons. Output metrics included total NeuN⁺, NeuN⁺ unedited, and NeuN⁺ edited cells, expressed as %NeuN⁺ edited to quantify neuronal editing efficiency. Control (RNP-only) samples were processed identically. Volumes were quantified by manually delineating the ROIs corresponding to the striatum across serial sections using FIJI (ImageJ). The total striatal volume was calculated based on the summed ROI areas multiplied by the distance between sections. Edited regions were identified by fluorescence (*e.g.*, tdTomato⁺ or GFP⁺ signal), and the edited volume was determined by summing the fluorescently positive ROIs and applying the same section spacing correction. Data were exported to spreadsheets and analyzed in GraphPad Prism (version 5.02; GraphPad Software, San Diego, CA, USA)

Image analysis for GFP editing and striatal volume with GFP loss was conducted using custom MATLAB scripts (MathWorks, Natick, MA). ROIs outlining each striatum were drawn on two-channel TIFF images (GFP, NeuN). NeuN⁺ neurons were segmented via adaptive contrast enhancement, Gaussian smoothing, adaptive thresholding, and connected-component labeling, optimized for neuronal morphology. GFP knockdown was quantified using a two-step thresholding of the inverted GFP channel: exclusion of bright pixels followed by Otsu’s method with a slight upward bias for specificity. Pixels above this threshold were labeled “GFP-off.” Neurons with ≥ 50% overlap between their NeuN mask and the GFP-off region were classified as GFP-off. GFP-off voxel counts were converted to physical volume (mm³). Output metrics included % GFP-off neurons and % GFP-off striatal area. Control samples were processed identically, and data were analyzed in GraphPad Prism (version 5.02; GraphPad Software, San Diego, CA, USA).

### NGS for m*HTT* genome editing

Genome editing was quantified by next generation sequencing (NGS) of PCR amplicons. Tissue was processed from multiple subsections using an Allprep DNA/RNA/Protein Mini Kit (Qiagen, CAT #80004), according to the manufacturer’s protocol. Genomic DNA was stored at −20°C. PCR amplification was performed using PrimeSTAR GXL DNA polymerase (Takara Bio) or Kapa HiFi (Roche) according to the manufacturer-provided protocol. Amplicons were purified using SPRI beads (UC Berkeley sequencing core), and concentrations were quantified via NanoDrop. Amplicon libraries were pooled and sequenced on an Illumina MiSeq at 300 bp paired end reads to a depth of at least 10, 000 reads per sample.

### Electroporation, plating, and genomic DNA isolation of R6/2 fibroblasts

R6/2 mice were humanely euthanized, and ears were diced and digested in collagenase D (Roche)–pronase (Neta Scientific) for 90 min at 37°C. Tissue was strained through a 70 µm filter, and cells were cultured in RPMI 1640 with 10% FCS, 50 µM 2-mercaptoethanol, 100 µM asparagine, 2 mM glutamine, and 1% penicillin-streptomycin. Primary fibroblasts (passage 3) were expanded in T75 flasks and electroporated (150, 000 cells/condition) using the P4 Primary Cell 96-well Nucleofector Kit and CA-137 program (Lonza). Post-electroporation, cells were split into two replicates per condition in 96-well plates. Genomic DNA was extracted ≥2 days post-editing with QuickExtract buffer (VWR) by incubating 10 min at room temperature, 20 min at 65°C, and 20 min at 95°C.

### DICOM Analysis for MRI reconstruction of putamen for large animal studies

Post-infusion T1-weighted MRI scans were co-registered with baseline images in Brainlab iPlan Flow Suite (v3.05). The putamen was manually delineated on baseline scans to generate a 3D ROI, and total volume was calculated in cm³. Anatomical landmarks were referenced using a pig brain stereotactic atlas. Post-infusion gadoteridol volumes were segmented using the SmartBrush tool, and infusion volumes were calculated. Coverage was defined as the percentage of putaminal volume containing gadoteridol.

### Genome safety test using targeted amplicon next generation sequencing

Tissues of interest were collected from treated animals and were flash-frozen in liquid nitrogen and stored at −80°C until DNA extraction. Genomic DNA was extracted using a modified phenol-chloroform protocol. Tissue samples were minced, placed into 2 mL screw-top tubes, and lysed overnight at 55°C with 580 µL Embryo Lysis Buffer and 20 µL Proteinase K (20 mg/mL) under rotation. Phenol-chloroform extraction was performed until the supernatant appeared clear, followed by a chloroform wash. DNA was precipitated by adding sodium acetate (final 10%) and 2× volume of ethanol, incubated at −80°C for 5 min, centrifuged, washed with 70% ethanol, air-dried, and resuspended in 200–300 µL 10 mM tris buffer. DNA concentration was measured using a NanoDrop spectrophotometer (Thermo Fisher, Cat # 13-400-5191P5). GFP on-target primers are in Table S6. Thermocycling conditions included an initial denaturation at 94°C for 1 min, 10 cycles of touchdown PCR (94°C, 30 s; 65–56°C, 30 s; 72°C, 30 s), followed by 25 cycles of standard PCR (94°C, 30 s; 56°C, 30 s; 72°C, 30 s), a final extension at 72°C for 5 min, and hold at 4°C. PCR products were purified via isopropanol precipitation and submitted to AZENTA Life Science (South Plainfield, NJ, USA) for paired-end sequencing, generating >100, 000 paired-end reads per sample. Sequencing data were analyzed using CRISPResso2 to detect gRNA editing in off-target tissues, including quality control.

### Open field test

Animals were placed in a 36 cm × 36 cm polypropylene open-field arena equipped with two rows of infrared sensors to detect horizontal and vertical movements (Open Field Photobeam Activity System, San Diego Instruments, Inc.). Arenas were housed in light- and sound-attenuating boxes. Activity counts were used to measure distance traveled, time resting, number of rears, and time spent in the center versus periphery.

### Rotarod test

Mice were tested over 2 consecutive days. On day 1, animals were habituated with three 1-min trials: the first with a stationary rod, the remaining at 16 RPM. Mice falling before 1 min were returned until habituated. On day 2, three accelerating trials (4–40 RPM) were conducted, recording latency to fall (max 5 min). Trials ended when mice fell or completed two rotations while clinging. Latencies were averaged for analysis.

### Statistics

Statistical analyses were performed using GraphPad Prism (version 5.02; GraphPad Software, San Diego, CA, USA). Data are presented as mean ± SD unless otherwise indicated. Comparisons among groups were conducted using one-way ANOVA followed by appropriate post hoc tests. Survival data were analyzed using Kaplan–Meier survival curves, with significance determined by log-rank (Mantel–Cox) and Gehan–Breslow–Wilcoxon tests.

## Acknowledgments

National Institute of Health, UG3 NS115599 (NM, KSB, RCW)

National Institute of Health, UH3 NS115599 (NM, KSB, RCW)

National Institute of Health, U19 NS132303 (CDC, KSB, RCW, RRL)

National Institute of Health, U42 OD026635 (SAM, CML)

National Institute of Health, U42 OD011140 (RSP, KL)

National Institute of Health, U42 OD027090 (KDW)

BDM was supported by the Curci Fellowship from the Shurl and Kay Curci Foundation. NM, KSB, and RCW were supported by UG3NS115599, UH3NS115599, and U19NS132303. RCW and NM are supported by Heritage Medical Research Institute. AJE is supported by the Kissick Family Foundation and the Shurl and Kay Curci Foundation. SAM is supported by U42 OD026635. RRL is supported by U19NS132303. CDC is supported by U19NS132303, the Carol and Gene Ludwig Foundation, the Rainwater Charitable Trust, and the Packard Foundation. We acknowledge the UC Berkeley DNA Sequencing Facility and The IGI Next Generation Sequencing Core (NGS Core) for their expertise. Confocal imaging experiments at UC-Berkeley were conducted at the CRL Molecular Imaging Center, RRID:SCR_017852, supported by UC Berkeley Biological Faculty Research Fund and Gordon and Betty Moore Foundation. We acknowledge Randy Walls and Jonathan Newall for their technical assistance for the validation study at JAX. We acknowledge Cathleen M. Lutz for her generous support in providing staff assistance for the validation study at JAX. Pigs (NSRRC:0016) used in this study were distributed by the National Swine Resource and Research Center (U42 OD011140). We further acknowledge support from the staff of the SCGE Swine Large Animal Testing Center (U42 OD027090) and the NextGen Precision Health MRI staff at the University of Missouri. We acknowledge Jennifer A. Doudna, Elizabeth Stahl, Dana Foss, Alzbeta Ressnerova, and Margaret Brown for their feedback and support in the development and review of this manuscript. We also acknowledge Sarah Pyle for assistance with graphical elements of the manuscript.

## Author contributions

Conceptualization: BDM, NM, PH, VSV, KSB, RCW

Methodology: BDM, MT, VM, CMB, LMAR, MHK, EAN, KA, RSC, EAN, NP, SKW, SHL, KJS, LS, JAG, KDW, AJE, NM, PH, VSV, KSB, RCW

Investigation: BDM, MT, VM, CMB, LMAR, MHK, RFH, EAN, KA, NP, RSC, RS, SKJW, DAK, NK, SHL, KJS, AKB, CSB, PAO, JB, AAO, MTR, MPZ, IG, GKS, MIP, EFS, ES, YG, ALS, PBS, LS, JAG, KDW, AJE, PH, VSV

Visualization: BDM, VM, CMB, LMAR, MHK, RSC, SHL, AKB, AJE, KJS, PH, VSV, RCW

Funding acquisition: NM, PH, VSV, KSB, RCW, KDW, SAM, CDC

Project administration: DAK, KJS, AKB, AAO, LS, JAG, KDW, SAM, NM, RRL, PH, VSV, KSB, RCW

Supervision: DAK, KJS, AKB, LS, JAG, KDW, SAM, AJE, CDC, NM, RRL, PH, VSV, KSB, RCW

Writing – original draft: BDM, MT, VM, CMB, KA, RSC, AKB, VSV, RCW

Writing – review & editing: BDM, MT, VM, CMB, KA, KJS, AKB, AJE, SAM, LS, CDC, VSV, RCW

## Data and materials availability

All data are available in the main text or the supplementary materials.

## Competing interests

RCW, LMAR, EAN, KA, and CMB are named inventors on one or more patent filings related to this work. RCW is a founder of Editpep, holds equity, and serves as an advisor. CMB holds equity in Editpep and serves as a consultant. NM is a founder of Opus Biosciences, Microbial Medical, and GenEdit. KSB holds equity in and/or financial positions in Editpep, Aviado Bio, Trogenix, and Precision NeuroMed. CDC is a founder with equity in Ciznor Co., a gene therapy company.

The other authors report no competing interests.

## Supplemental Figures

**S1:**
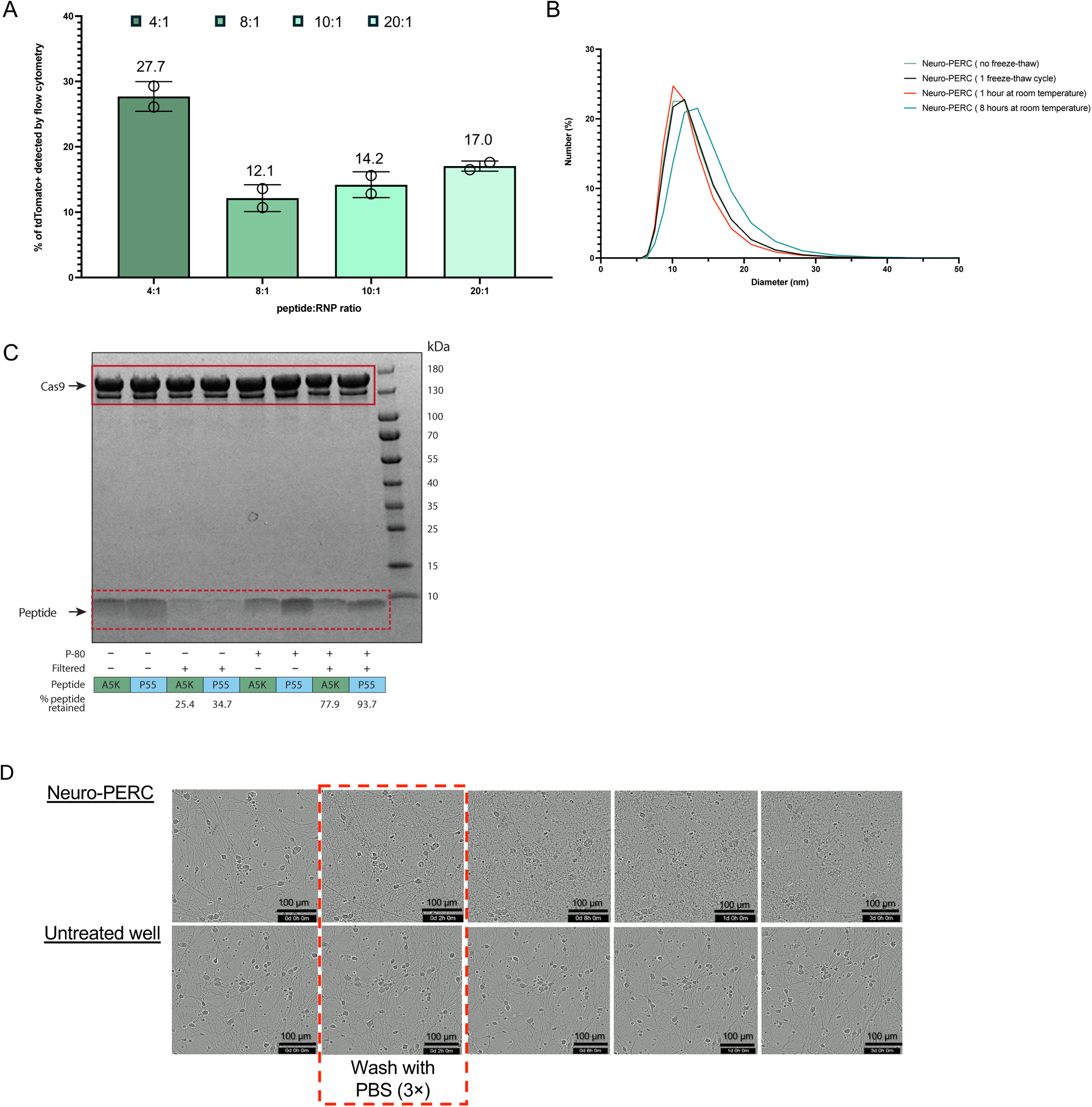
Optimization of Neuro-PERC formulation *ex vivo*. **(A)** Quantification via flow cytometry of murine tdTomato^+^ neural progenitor cells (NPCs), indicating successful editing across formulations with different peptide:RNP ratios. Each data point represents one biological replicate (n = 2). Data are shown as mean ± SD **(B)** DLS analysis showing size distributions of Neuro-PERC formulations by number. The optimized formulation forms monodisperse nanoparticles (∼17 nm diameter) that remain stable after one freeze–thaw cycle and up to 8 h at RT **(C)** SDS–PAGE confirming peptide retention in Neuro-PERC with A5K and P55, demonstrating enhanced retention after filtration (25.4–34.7% unfiltered; 77.9–93.7% filtered). kDa, kilodalton. Solid red line, Cas9; dotted red line, peptide. **(D**) Post-mitotic human iPSC-derived neurons were treated with Neuro-PERC formulations targeting B2M at 0 h (start of time course). A triple Dulbecco’s Phosphate-Buffered Saline (D-PBS) wash was performed at 2 h (indicated by the red dashed line). Scale bar, 100 µm.

**S2:**
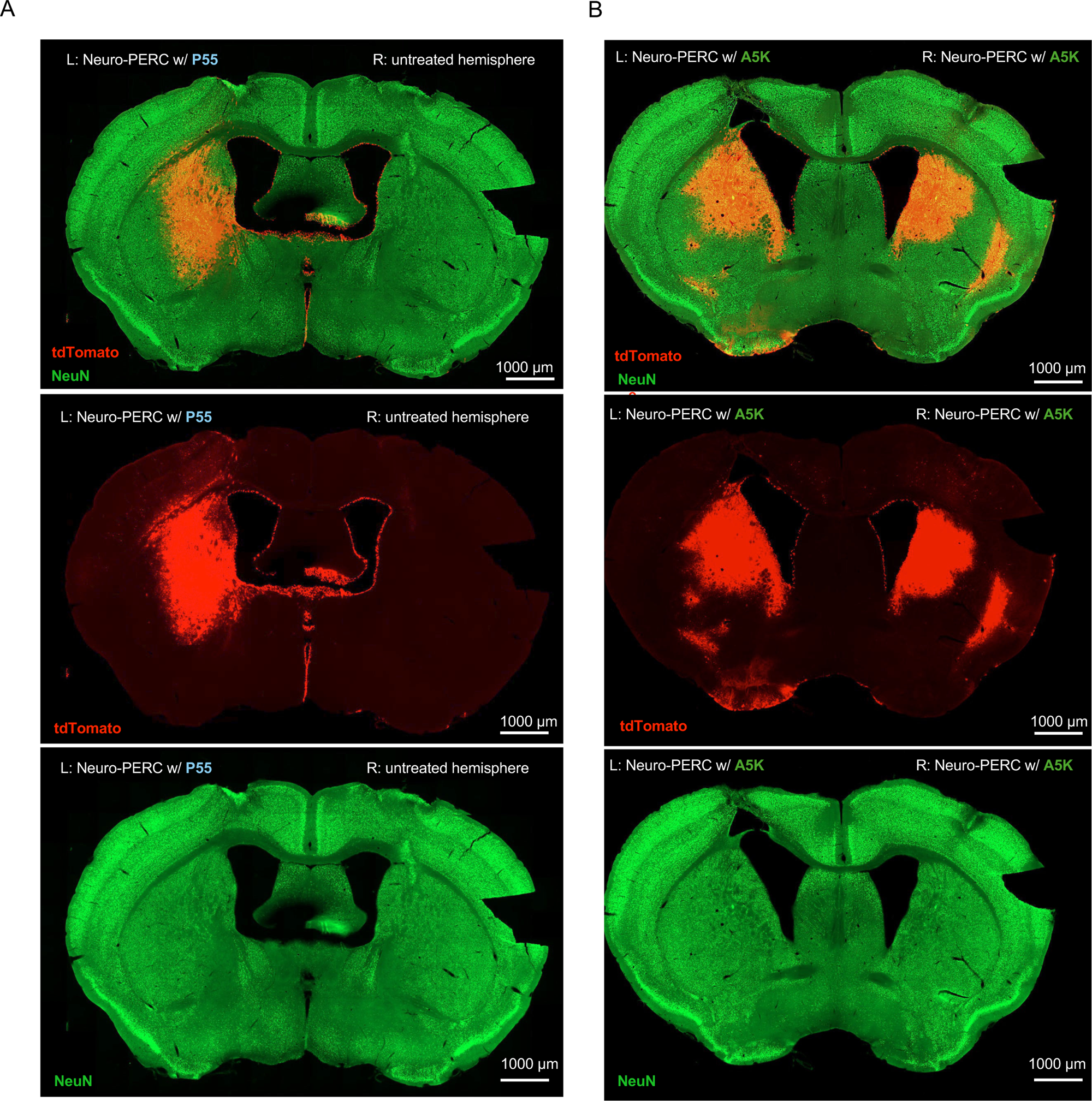
Split-channel visualization of Ai9 mouse brains following Neuro-PERC administration via CED. **(A)** Merged and individual channels showing NeuN (green) and tdTomato (red) in P55-based NeuroPERC treated and untreated hemispheres. Scale bar: 1000 µm **(B)** Merged and individual channels showing NeuN (green) and tdTomato (red) in A5K-based NeuroPERC treated hemispheres. Scale bar: 1000 µm.

**S3:**
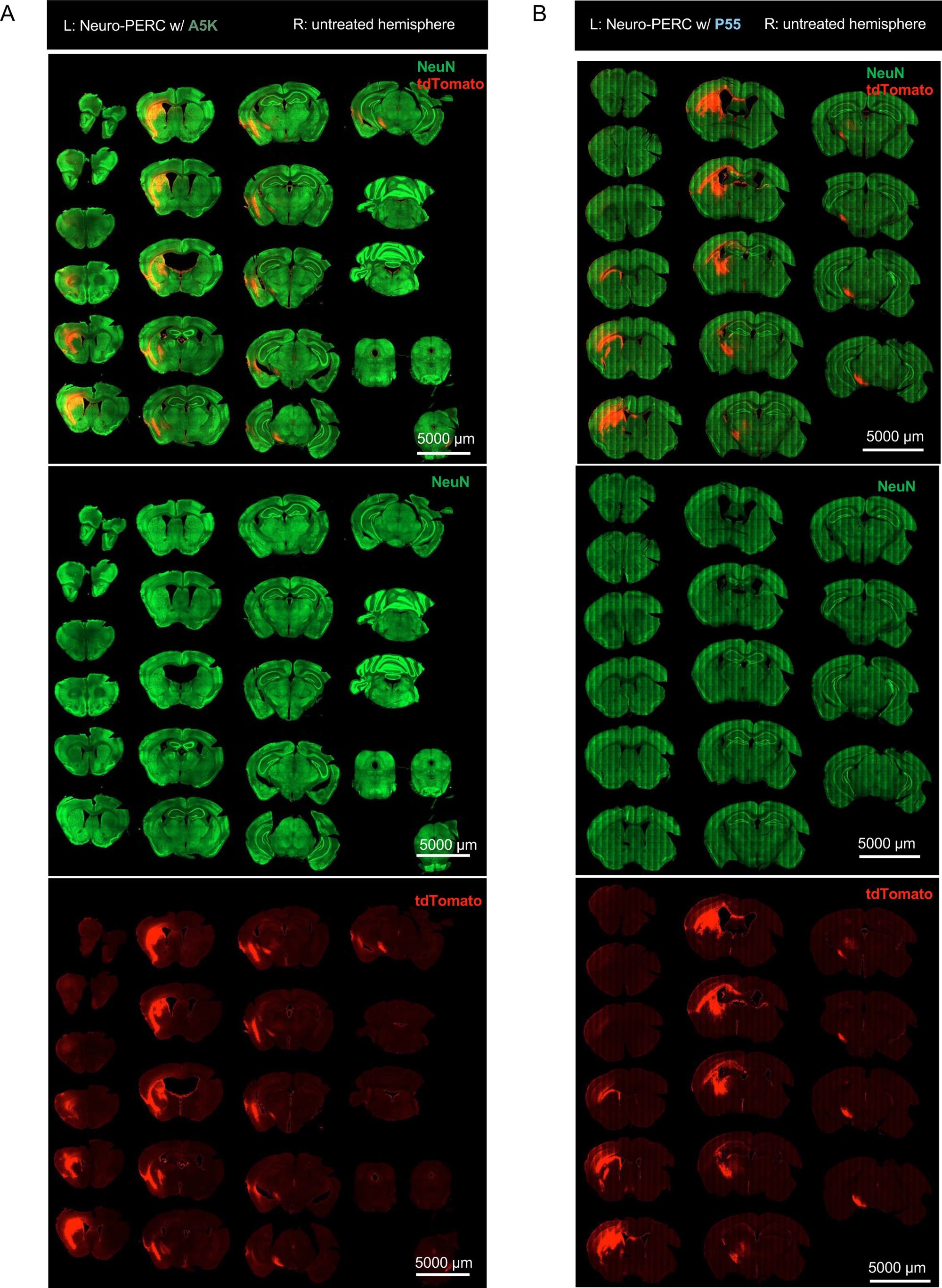
Whole-brain serial coronal sections of Ai9 mouse brains following Neuro-PERC administration via CED. **(A)** Serial coronal sections from a representative mouse treated with A5K-based Neuro-PERC, showing NeuN (green), tdTomato (red), and merged channels across the entire brain. Left hemisphere = treated; right = untreated. Scale bar: 5000 µm (**B)** Serial coronal sections from a mouse treated with P55-based Neuro-PERC, showing NeuN (green), tdTomato (red), and merged channels across the entire brain. Left hemisphere: treated; right: untreated. Scale bar: 5000 µm.

**S4:**
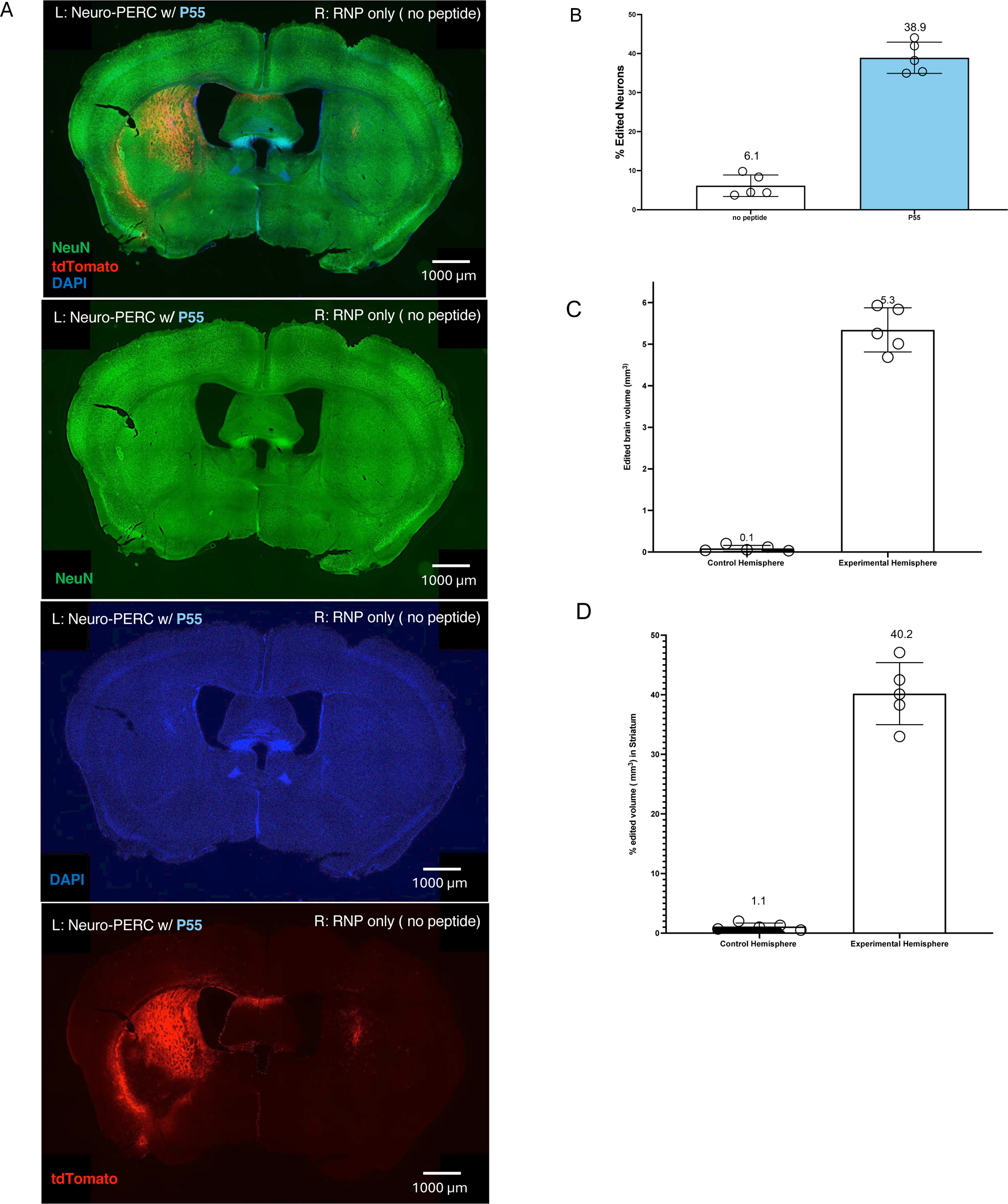
Independent validation of P55-based Neuro-PERC mediated genome editing in Ai9 mice performed at the Jackson Laboratory (JAX) (A) Representative immunofluorescence sections show robust tdTomato expression (red) in the P55-based Neuro-PERC treated hemisphere (left), with NeuN (green) and DAPI (blue). Scale bar: 1000 µm (B–D) Quantitative analyses for the treated and untreated hemispheres showing (B) the percentage of tdTomato⁺/NeuN⁺ neurons (C) edited area normalized to striatum, and (D) total edited volume as a proportion of striatal volume. Each dot represents one biological replicate (n = 5).

**S5:**
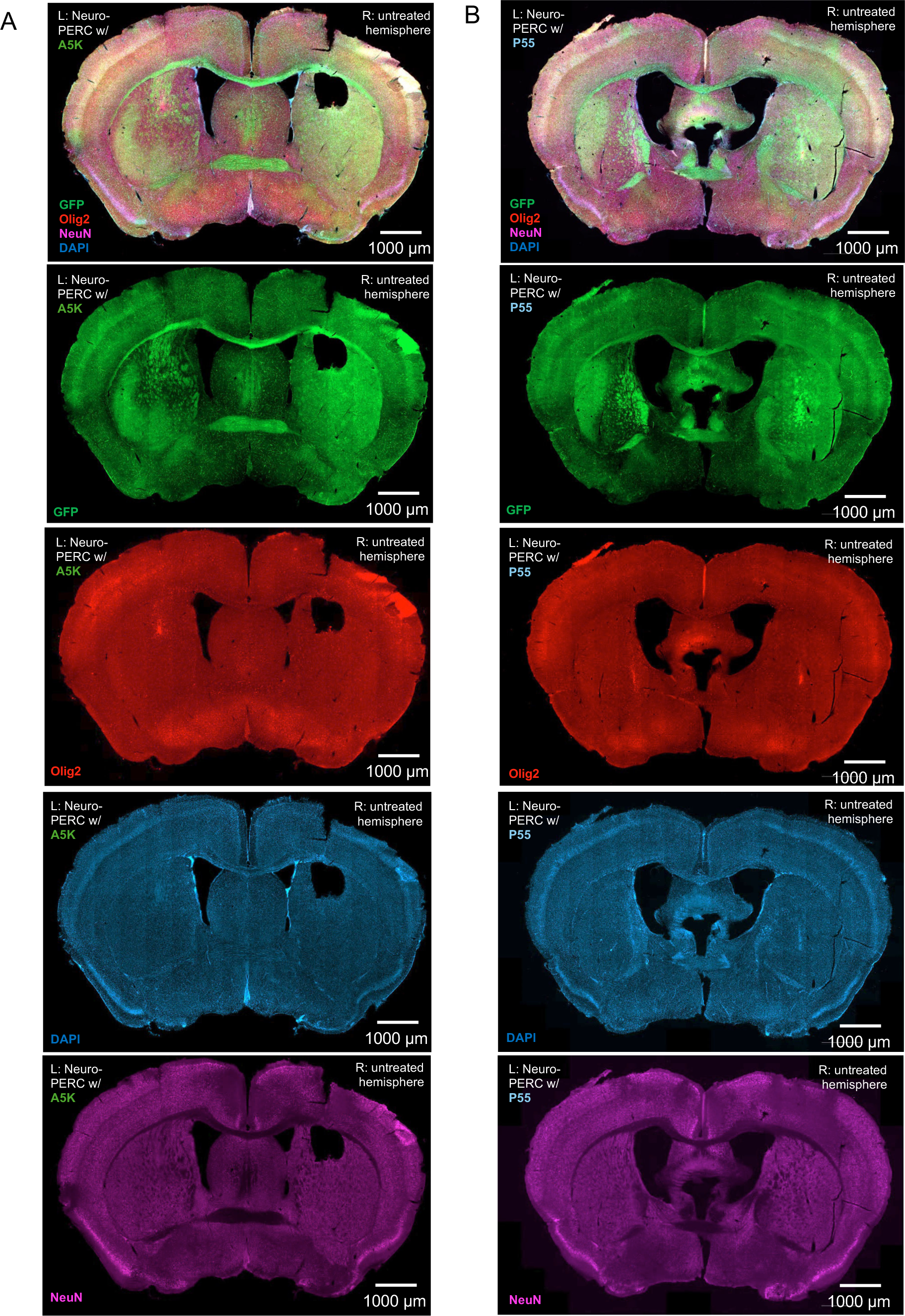
Split-channel visualization of GFP mouse brains following *in vivo* Neuro-PERC administration via CED. **(A)** Merged and individual channels showing GFP (green), Olig2 (red), NeuN (magenta), and DAPI (blue) in A5K-based Neuro-PERC–treated and untreated hemispheres. Scale bar, 1000 µm **(B)** Merged and individual channels showing GFP (green), Olig2 (red), NeuN (magenta), and DAPI (blue) in P55-based Neuro-PERC–treated and untreated hemispheres. Scale bar, 1000 µm.

**S6:**
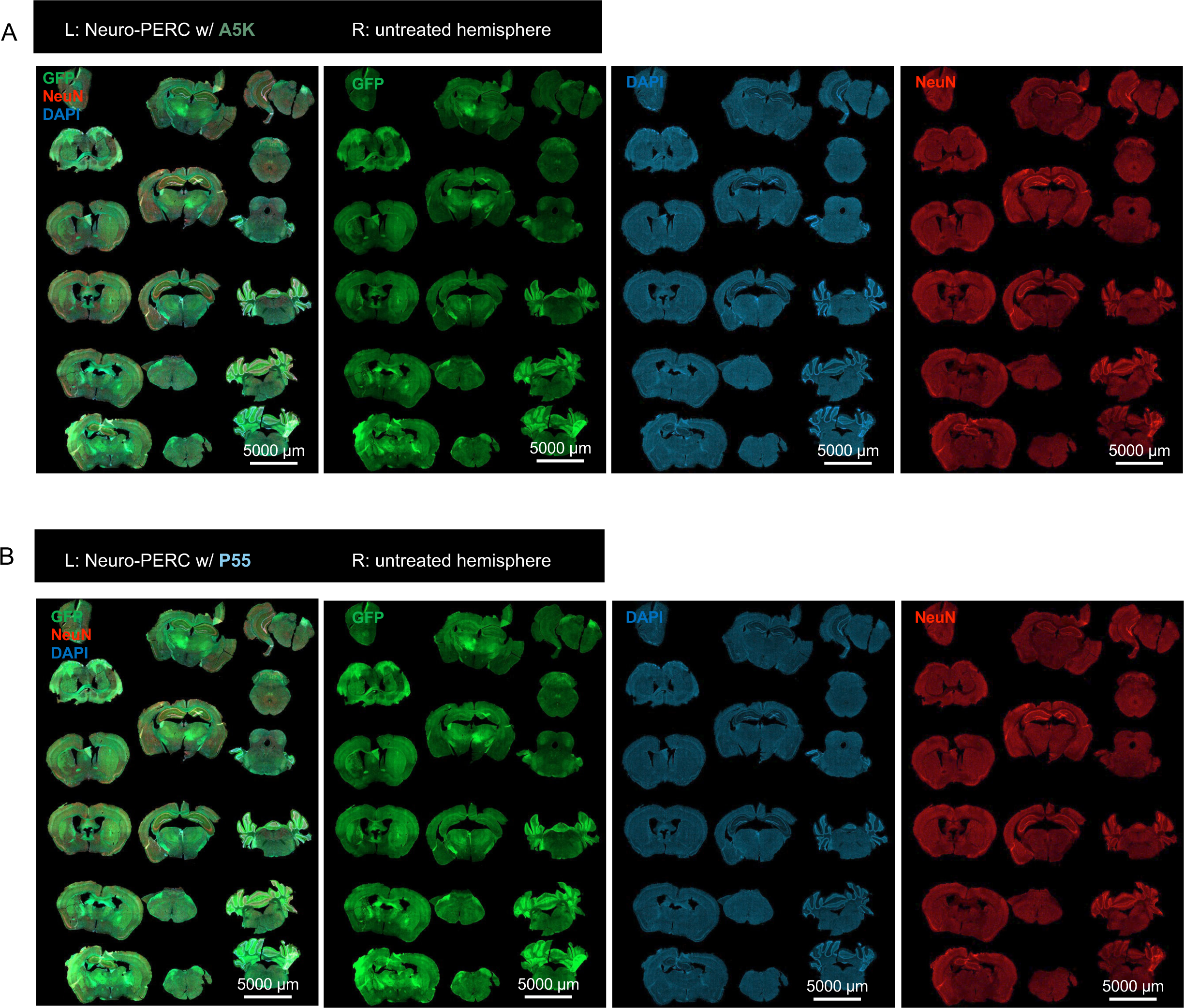
Whole-brain serial coronal sections of GFP mouse brains following Neuro-PERC administration via CED. **(A)** Serial coronal sections from a mouse treated with A5K-based Neuro-PERC, showing GFP (green), NeuN (red), DAPI (blue), and merged channels across the entire brain. Left hemisphere: treated; right: untreated. Scale bar, 5000 µm **(B)** Serial coronal sections from a mouse treated with P55-based Neuro-PERC, showing GFP (green), NeuN (red), DAPI (blue), and merged channels across the entire brain. Left hemisphere = treated; right = untreated. Scale bar, 5000 µm.

**S7:**
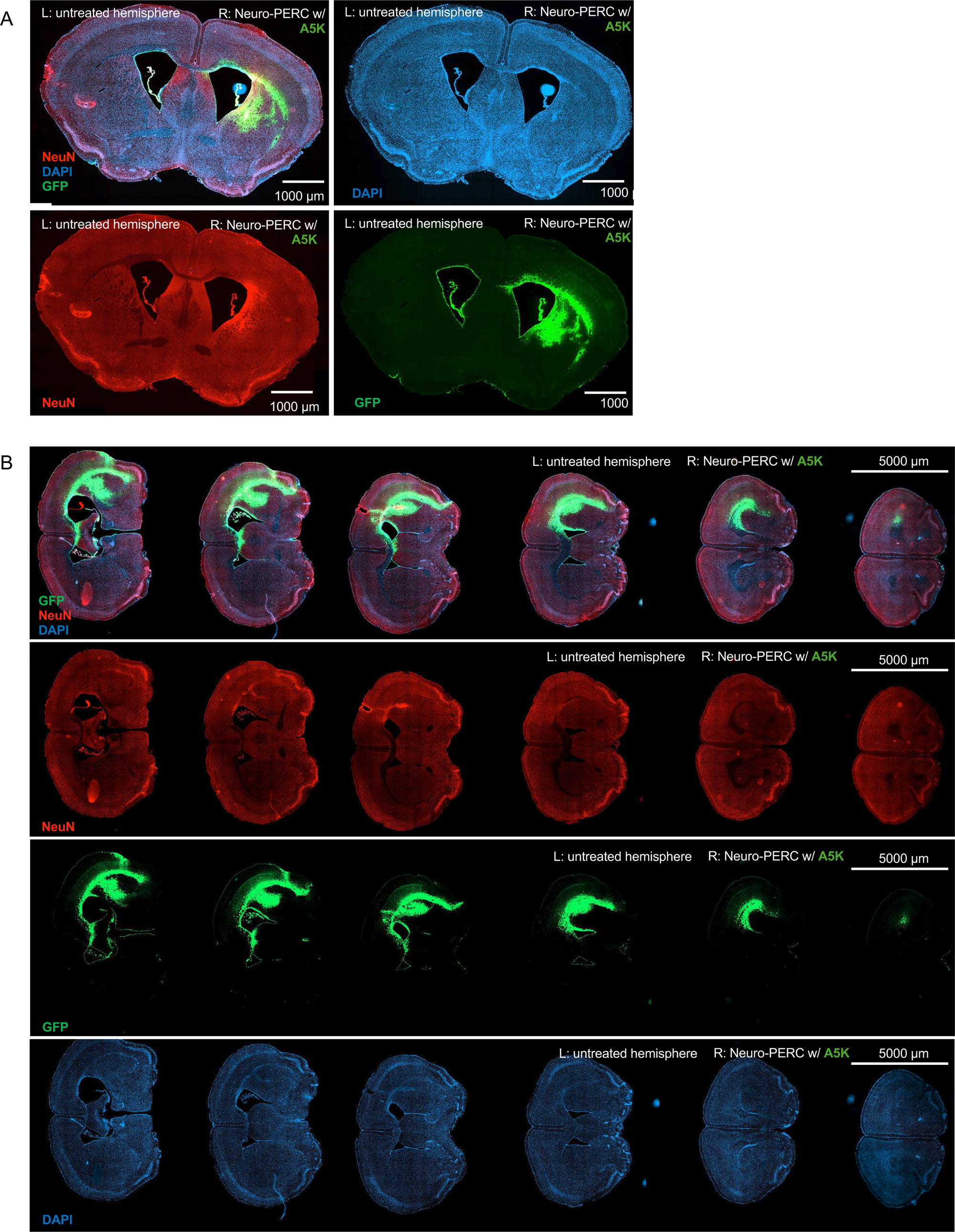
*In vivo* adenine base editing in GFP reporter mice (GER10) following administration of ABE-8e/A5K Neuro-PERC with CED. **(A)** Merged and individual channels showing GFP (green), NeuN (red), and DAPI (blue) in Ger10 mice treated with ABE-8e/A5K Neuro-PERC (right hemisphere) and the untreated hemisphere (left). Scale bar, 1000 µm. **(B)** Serial sections from the same mouse treated ABE-8e/A5K Neuro-PERC with showing merged and individual channels for GFP (green), NeuN (red), and DAPI (blue) across the entire brain. Left hemisphere: untreated; right ABE-8e/A5K Neuro-PERC.

**S8:**
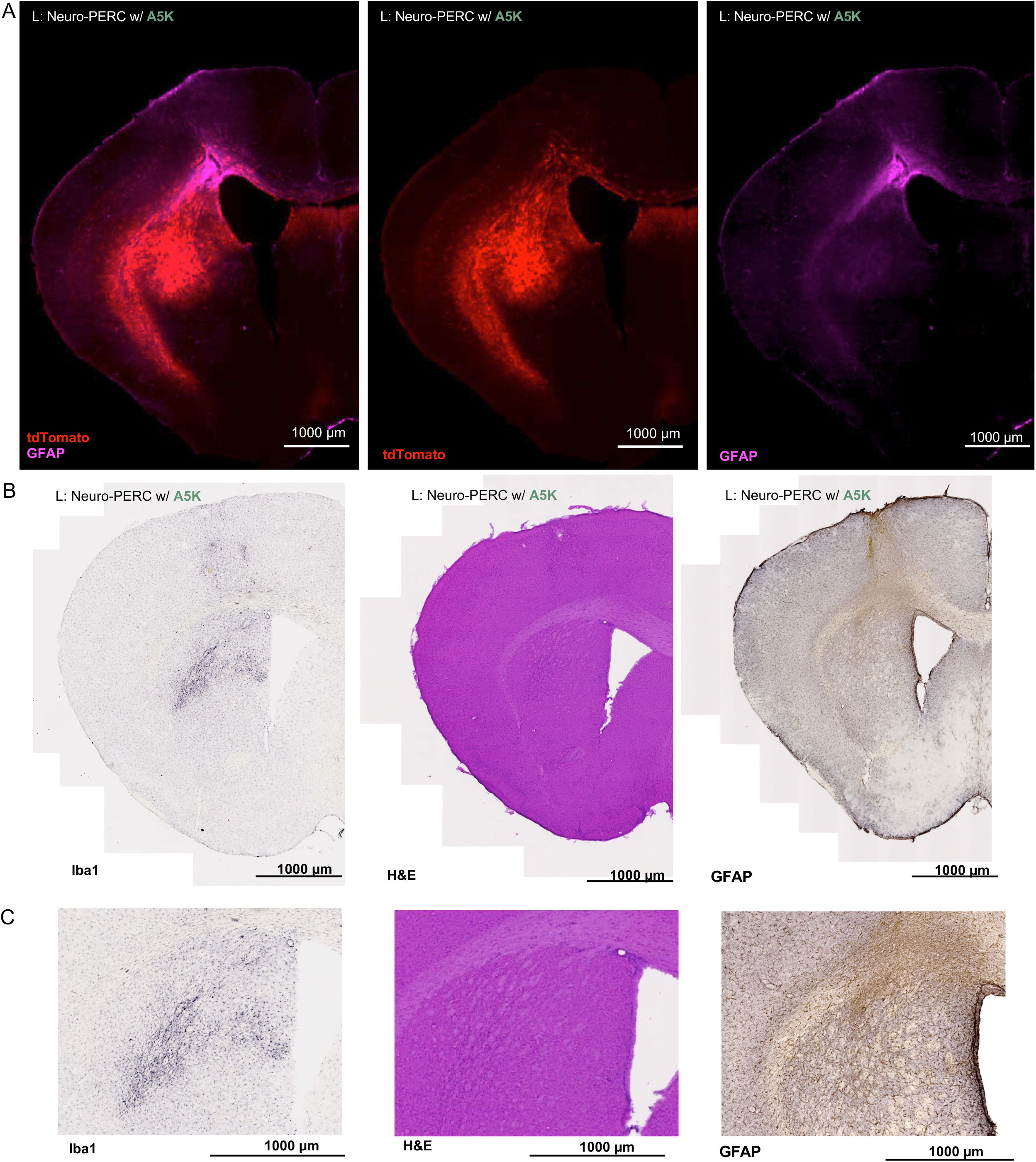
Evaluation of local immune response in Ai9 mice at 3 weeks after the administration of A5K-based Neuro-PERC with CED. **(A)** Merged and individual fluorescence channels showing tdTomato (red) and GFAP (purple) in a representative brain section from an Ai9 mice treated with A5K-based Neuro-PERC (left hemisphere). Scale bar, 1000 µm **(B)** Adjacent sections processed using 3, 3′-diaminobenzidine (DAB) -based chromogenic staining for Iba1 (microglia, gray) and GFAP (astrocytes, brown), along with hematoxylin-and-eosin (H&E, purple) staining to assess tissue morphology. Scale bar, 1000 µm **(C)** Higher-magnification DAB images from panel B showing detailed cellular morphology in regions adjacent to the infusion site for Iba1, H&E, and GFAP. Scale bar, 1000 µm.

**S9:**
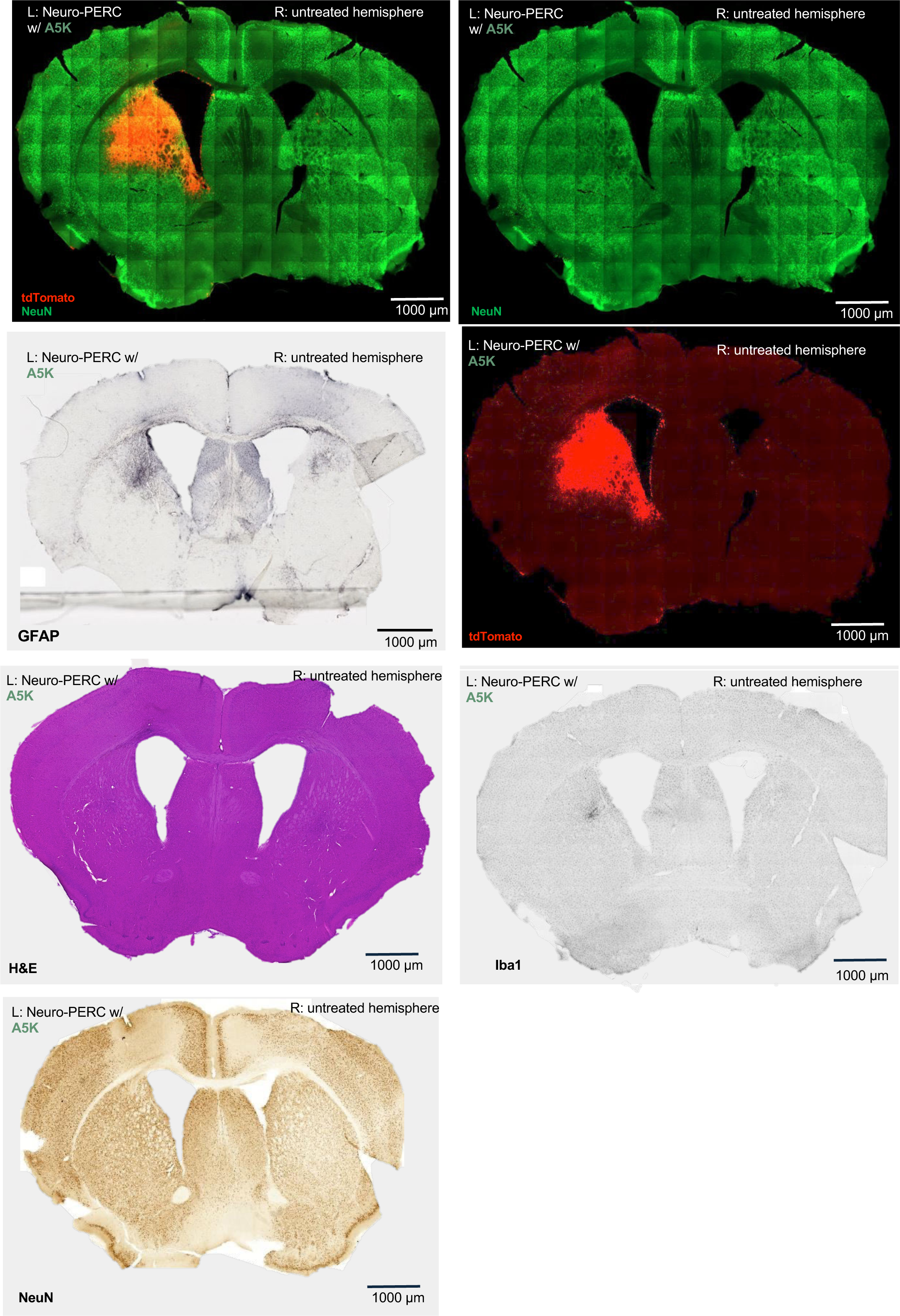
Evaluation of local immune response in Ai9 mice at 3 months after the administration of A5K-based Neuro-PERC with CED. Representative brain sections from an Ai9 mice 3 months post-administration showing tdTomato (red) and NeuN (green) fluorescence, along with adjacent sections stained for GFAP (astrocytes, gray), Iba1 (microglia, gray), hematoxylin and eosin (H&E; purple), and NeuN (brown, DAB-based). Left hemisphere A5K-based Neuro-PERC: right: untreated hemisphere. Scale bars, 1000 µm.

**S10:**
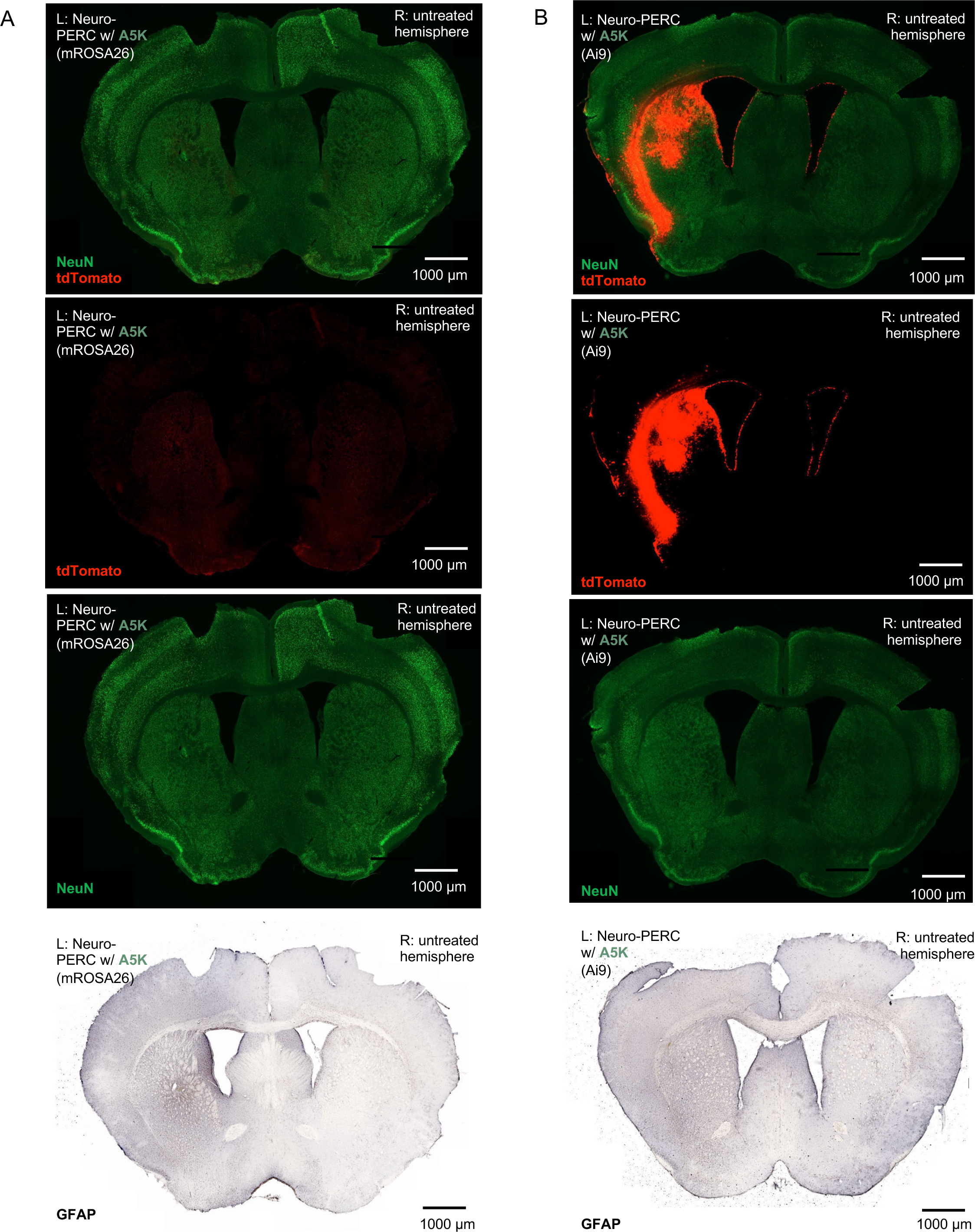
Comparison of local immune response at 3 weeks following administration of A5K-based Neuro-PERC with mROSA26 or Ai9 guide RNPs. **(A)** Representative brain sections from Ai9 mice treated with A5K-based Neuro-PERC carrying the non-targeting mROSA26 guide (left hemisphere) and buffer control (right hemisphere). Sections were immunostained for NeuN (green), tdTomato (red), and GFAP (gray, astrocyte marker). Scale bar, 1000 µm **(B)** Representative brain sections from Ai9 mice treated with A5K-based Neuro-PERC carrying the non-targeting Ai9 guides (left hemisphere) and buffer control (right hemisphere). Sections were immunostained for NeuN (green), tdTomato (red), and GFAP (gray, astrocyte marker). Scale bar, 1000 µm.

**S11:**
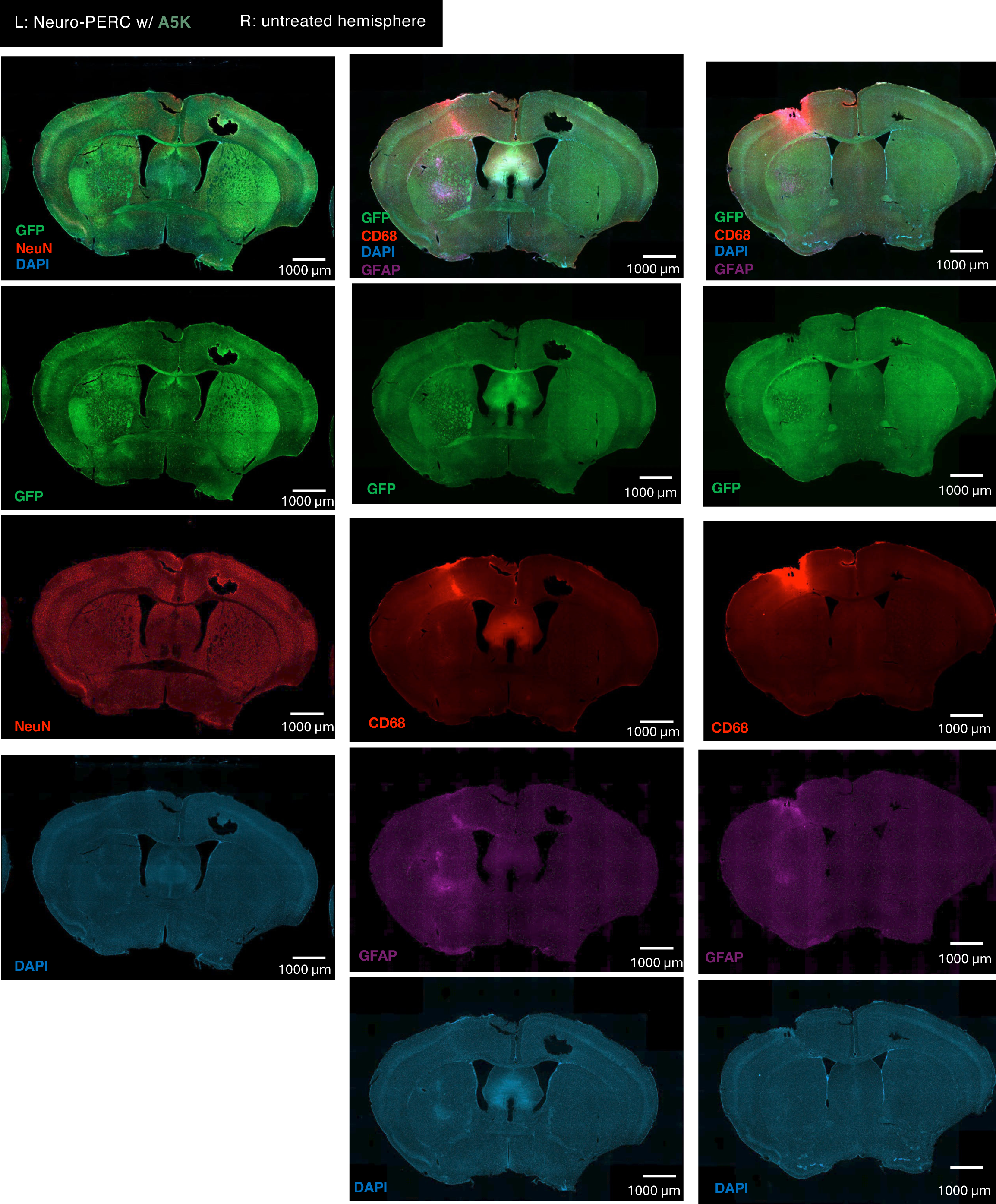
Evaluation of local immune response and microglial activation in GFP mice at 3 weeks after the administration of A5K-based Neuro-PERC with CED. Left panel: merged and individual fluorescence channels showing GFP (green), NeuN (red), and DAPI (blue) in coronal brain sections from GFP reporter mice treated with A5K-based Neuro-PERC (left hemisphere) and the untreated hemisphere (right). Scale bar, 1000 µm Right and Middle panels: merged and individual fluorescence channels showing GFP (green), CD68 (red), GFAP (magenta), and DAPI (blue) in coronal brain sections from the same animals. Scale bar, 1000 µm.

**S12:**
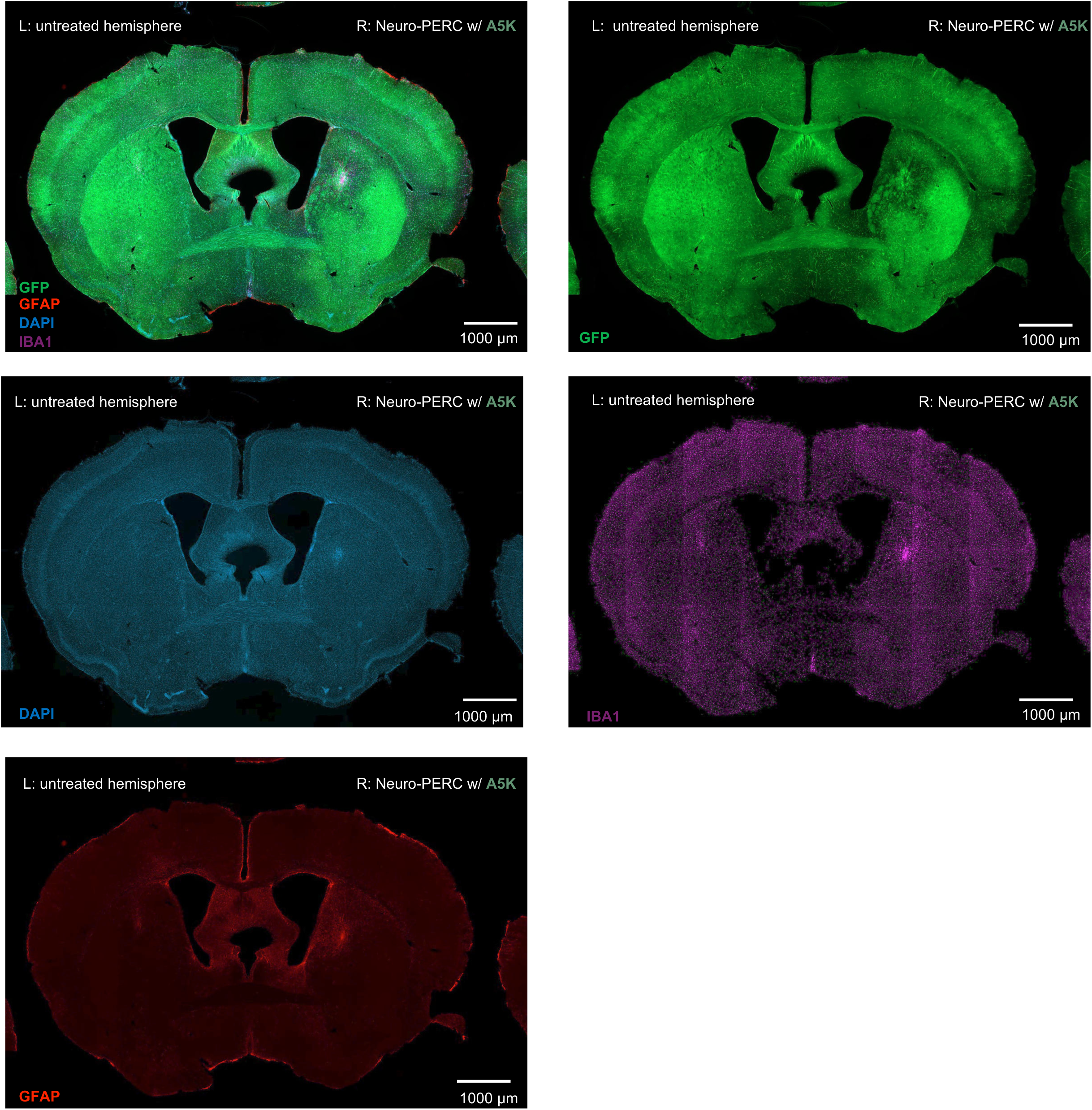
Evaluation of local immune response and microglial presence in GFP mice at 3 months after the administration of A5K-based Neuro-PERC with CED. Representative brain sections from GFP reporter mice 3 months post-administration showing GFP (green) fluorescence, along with adjacent sections stained for GFAP (astrocytes, red), Iba1 (microglia, magenta), and DAPI (blue). Right hemisphere: A5K-based Neuro-PERC; Left: untreated hemisphere. Scale bars, 1000 µm.

**S13:**
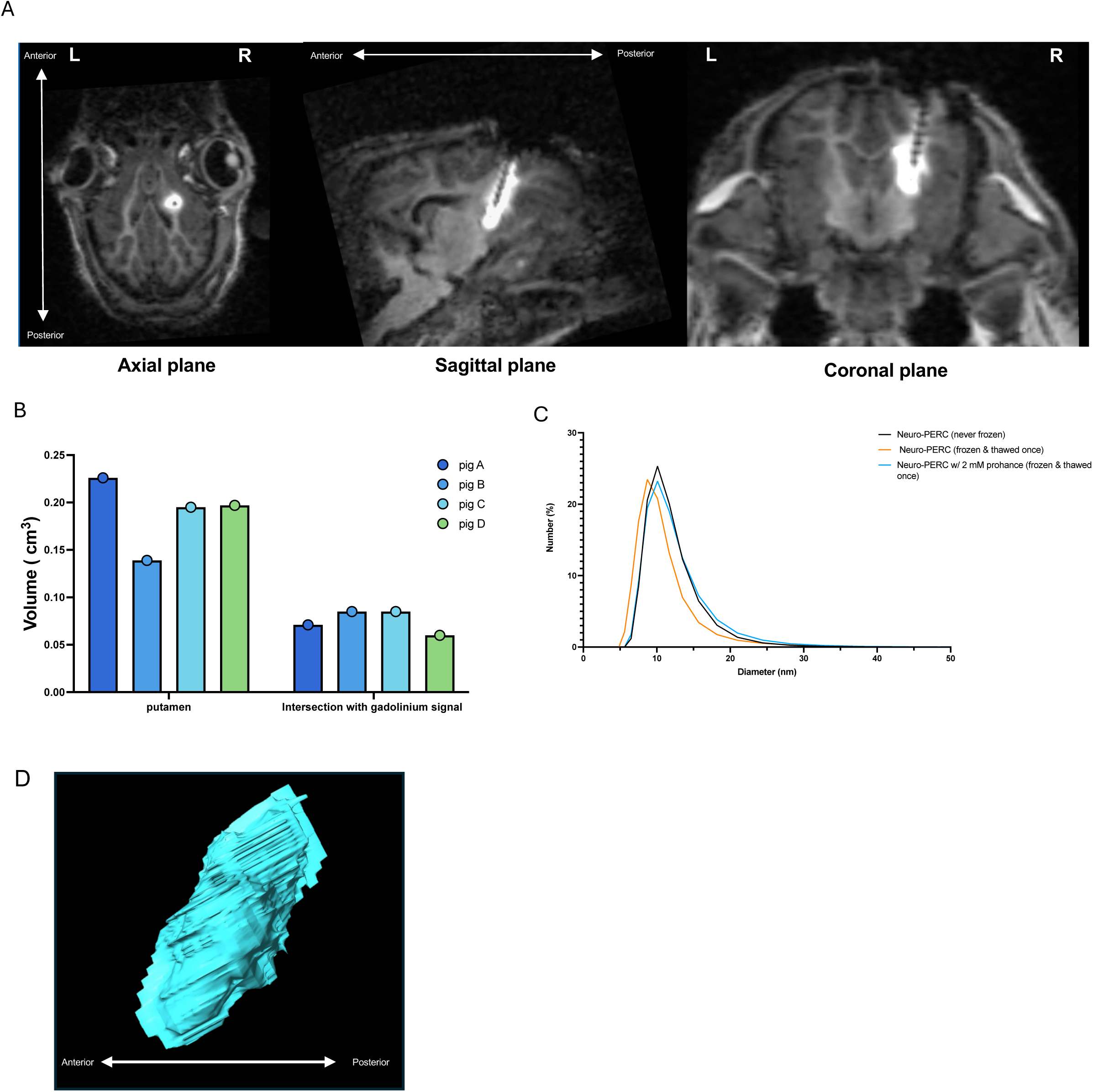
MRI-guided convection-enhanced delivery (CED) and formulation stability of Neuro-PERC in pigs. **(A)** Representative axial, sagittal, and coronal MRI planes acquired during real-time, gadolinium-enhanced convection-enhanced delivery (CED) of A5K-based Neuro-PERC into the putamen of the pig brain. Gadolinium (ProHance, 2 mM) was co-infused to enable visualization of the infusion cloud and verification of cannula placement, infusion spread, and target coverage within the putamen. The bright white signal is the ProHance (**B)** Quantification of infused putaminal volume and overlap with the gadolinium signal, illustrating infusion reproducibility across biological replicates (n = 4 pigs) **(C)** DLS Showing particle-size distributions of Neuro-PERC under different handling conditions: freshly prepared, after one freeze–thaw cycle, and following post–freeze thaw with 2 mM ProHance (comparable to surgical conditions). **(D)** A 3D reconstruction of the infusate was generated from the post-infusion DICOM images using the Brainlab iPlan Flow Suite (v3.05) and its SmartBrush semiautomated segmentation tools in the axial, sagittal, and coronal planes.

**S14:**
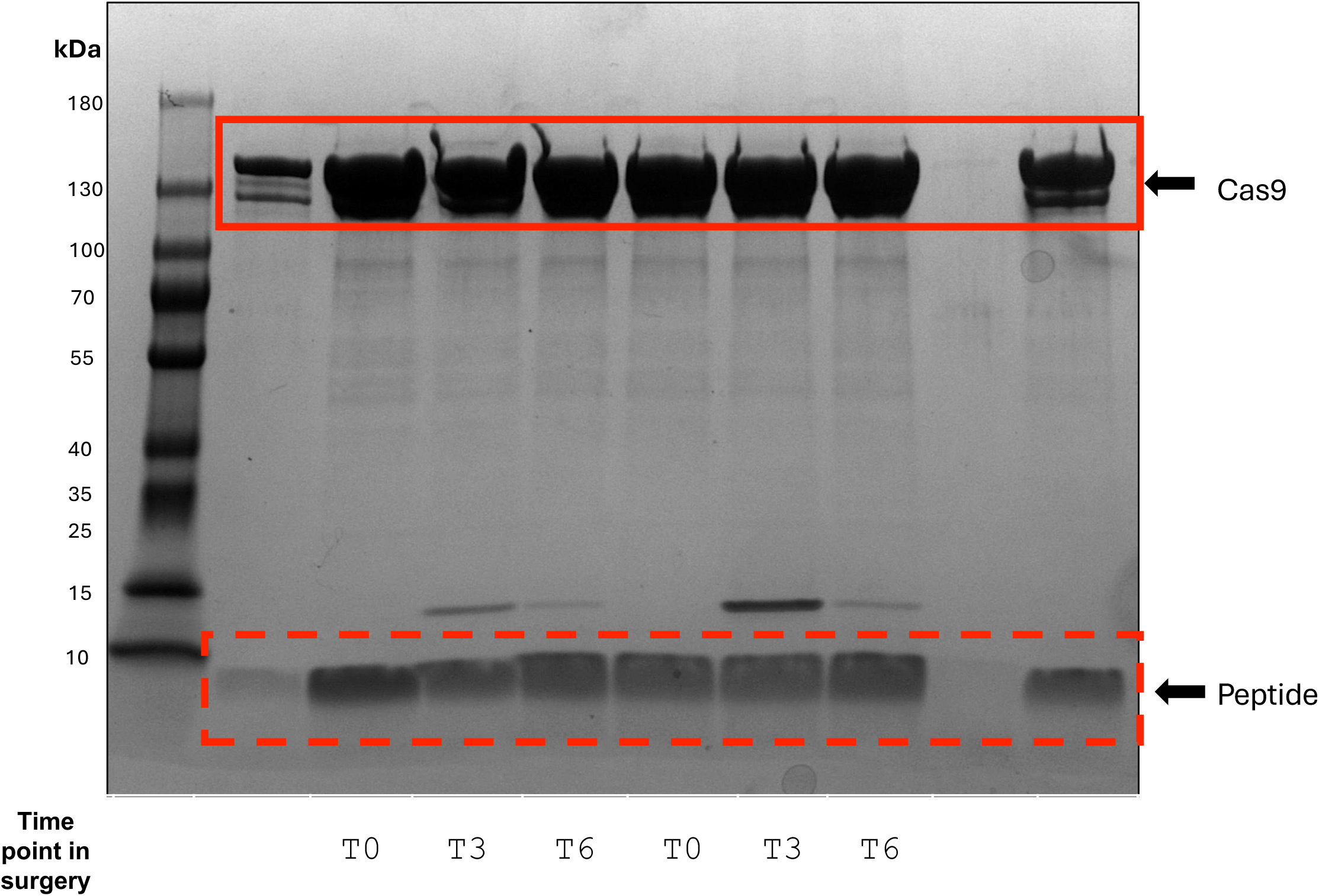
SDS–PAGE analysis of Neuro-PERC material collected at various intraoperative time points during the pig surgeries across 2 days. Samples were analyzed from the infusion line during MRI-guided convection-enhanced delivery (CED) of Neuro-PERC spiked with ProHance contrast agent. T0 represents the material collected immediately before infusion (freshly spiked), T3 corresponds to 3 hours into the procedure (during pig switch), and T6 marks completion of infusion (for 2 pigs). The first lane contains a 1:10 dilution of the formulation, while the second T0–T6 series reflects the next-day surgical preparation. The second-to-last lane shows the priming solution containing peptide for the surgical tubing, and the final lane represents the formulation control. Prominent bands correspond to Cas9 and peptide A5K.

**S15:**
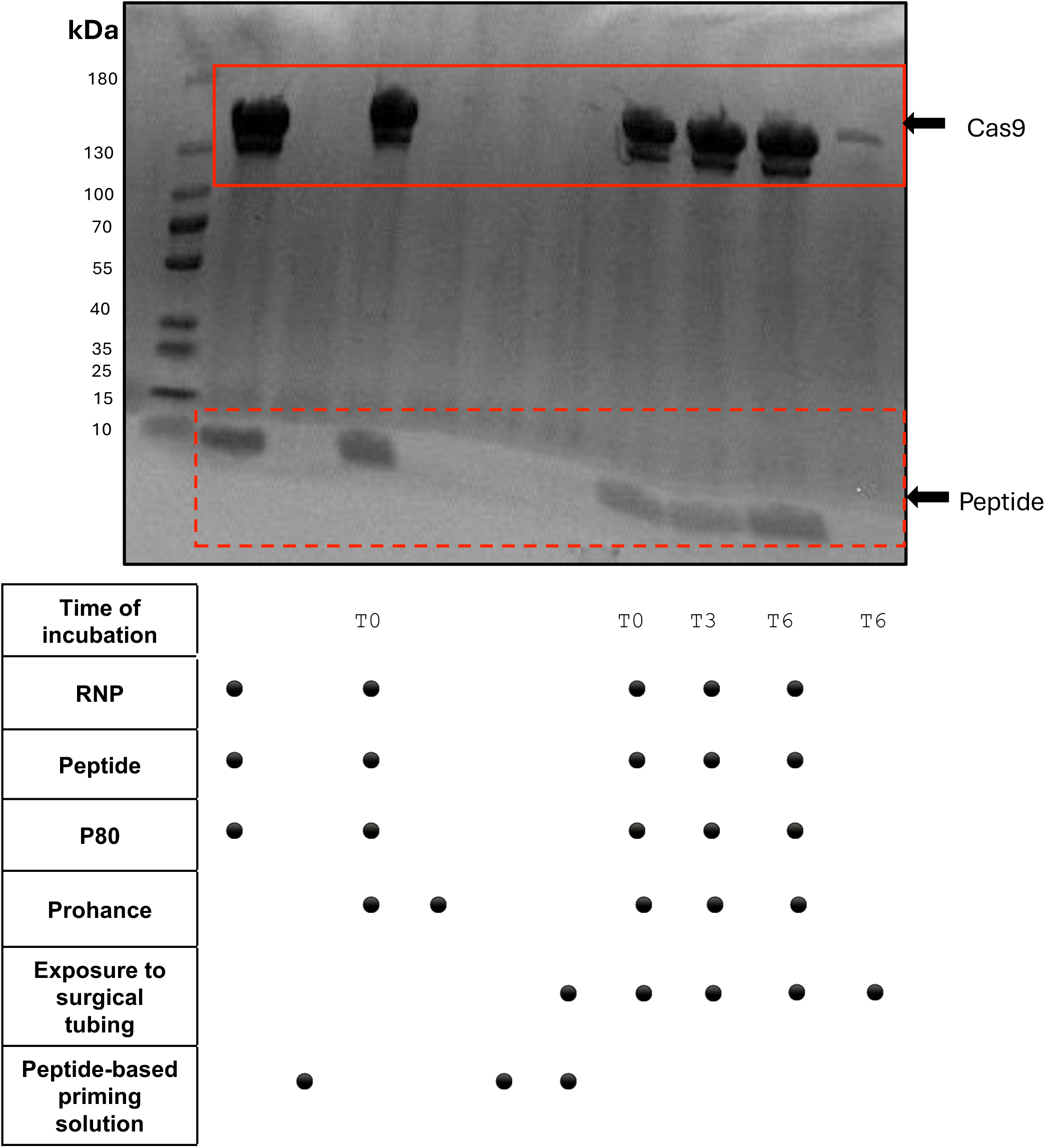
*Ex vivo* simulation of intraoperative Neuro-PERC to assess peptide retention following exposure to surgical tubing. SDS–PAGE analysis of Neuro-PERC formulations which simulated the intraoperative workflow. Samples of Neuro-PERC. The samples were prepared with or without ProHance and incubated for defined time points (T0, T3, T6) to mimic the duration of surgical handling in the surgical tubing. A peptide based priming solution with P-80 was used for line pre-fill prior to administration to limit the loss of material during the procedure. The matrix below summarizes formulation components and exposure conditions for each lane. prominent bands correspond to Cas9 and peptide A5K.

**S16:**
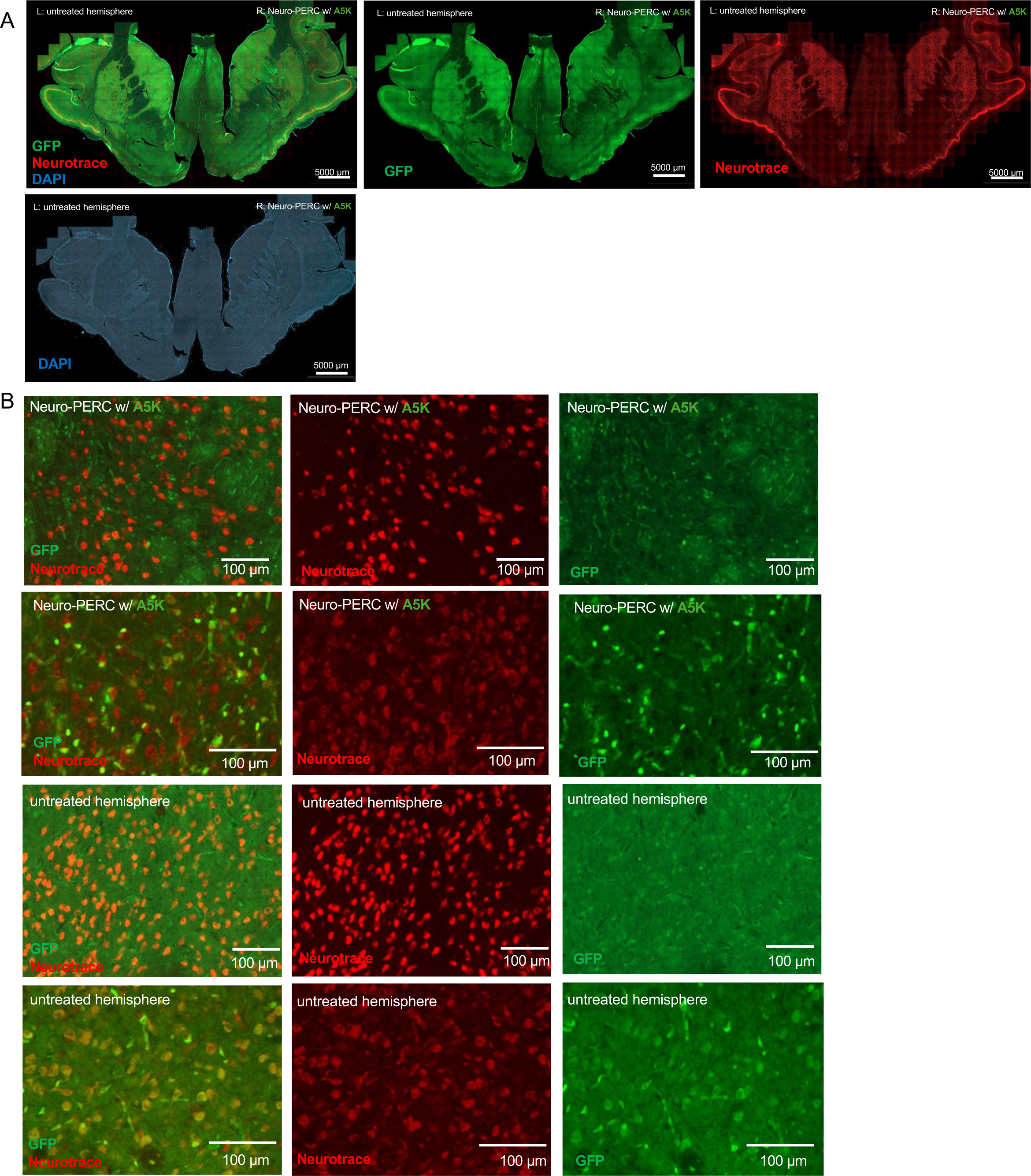
Split-channel visualization of pig brain sections following A5K-based Neuro-PERC administration via MRI-guided CED at 1-month post-surgery. **(A)** Merged and individual channels showing GFP (green), Neurotrace (red), and DAPI (blue) in coronal brain sections from the pig infused with A5K-based Neuro-PERC (right hemisphere) and the untreated hemisphere (left) with MRI-guided CED. Scale bar, 5000 µm Merged and individual channels of higher magnification microscopy showing GFP (green), Neurotrace (red) in coronal brain sections from the pig infused with A5K-based Neuro-PERC (top 2 rows) and an untreated hemisphere as a control (bottom 2 rows);. Scale bar, 100 µm.

**S17:**
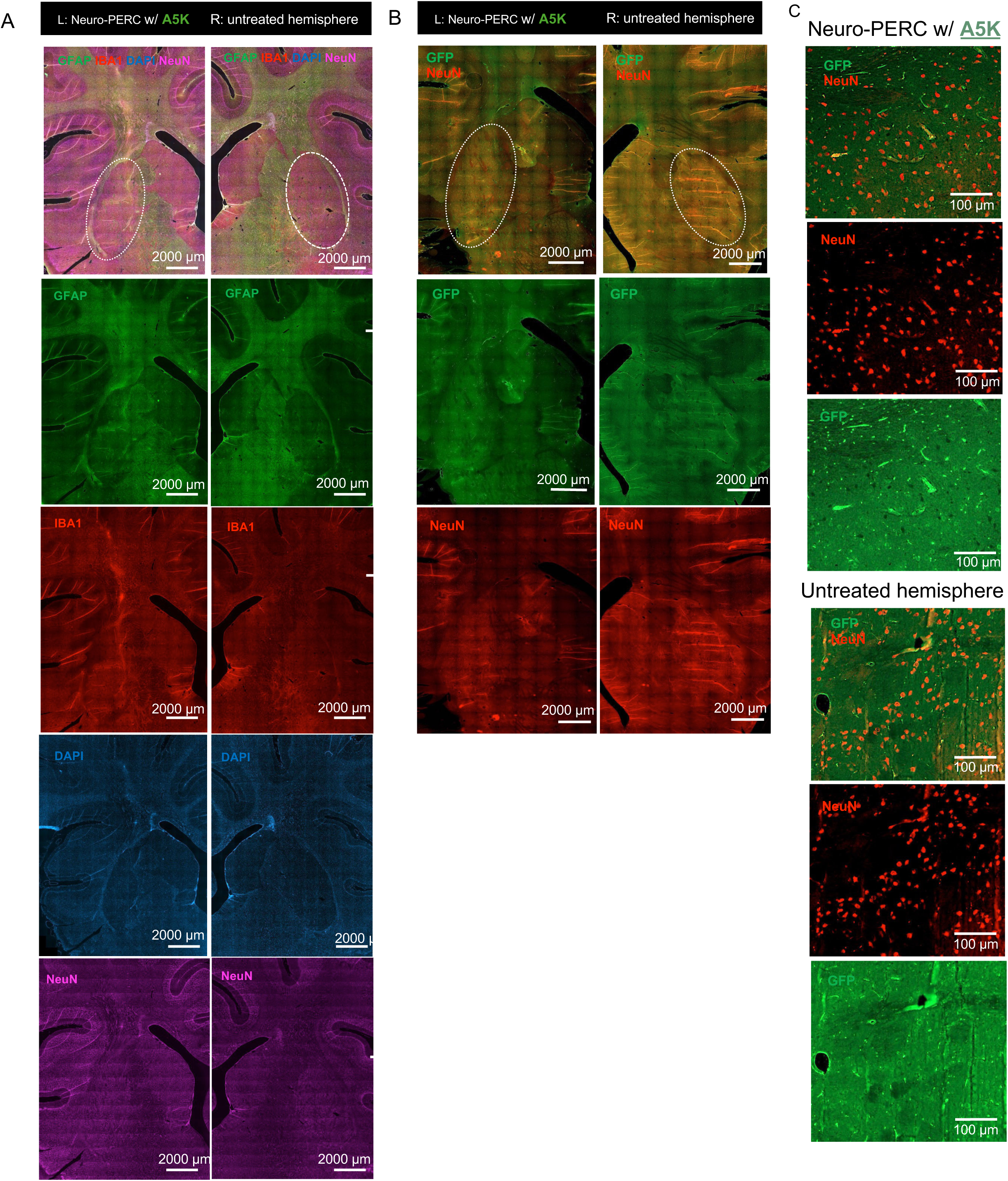
Split-channel visualization of pig brain sections 3 months after A5K-based Neuro-PERC administration via MRI-guided CED. **(A)** Merged and individual fluorescence channels showing GFAP (green), Iba1 (red), NeuN (magenta), and DAPI (blue) in the left hemisphere (A5K-based Neuro-PERC–treated) and right hemisphere (untreated). The targeted region (putamen): white dotted line; Scale bar, 2000 µm **(B)** Merged and individual channels show GFP (green) and NeuN (red) from the same animal. The targeted region (putamen): white dotted line; Scale bar, 2000 µm (**C)** High magnification of targeted region of the left hemisphere (A5K-based Neuro-PERC–treated) and right hemisphere (untreated); Scale bar, 100 µm.

**S18:**
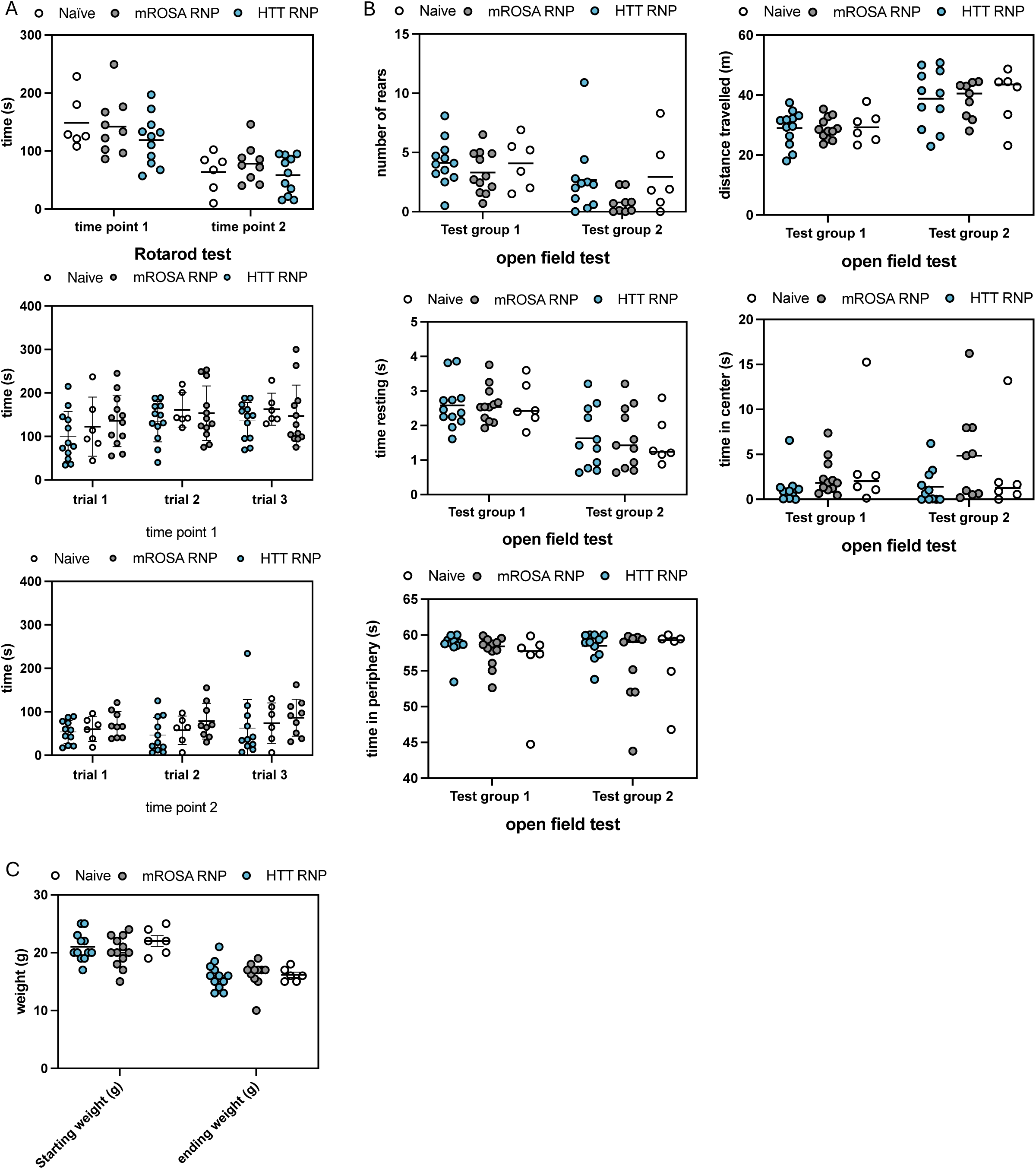
Behavioral assessment of R6/2 mice treated with A5K-based Neuro-PERC carrying HTT-targeting or control RNPs. **(A)** Rotarod performance of R6/2 mice treated with Neuro-PERC carrying HTT-targeting RNP (*blue*), mROSA26 RNP (*gray*), or untreated naïve (*white*) controls. Graphs show latency to fall (time on rotarod) measured across three trials at two independent time points. Each dot represents one mouse. **(B)** Open-field behavioral analysis showing number of rears, time spent resting, distance traveled, time in center, and time in periphery for the same treatment groups — HTT RNP (*blue*), mROSA26 RNP (*gray*), and naïve (*white*) (**C)** Starting and ending body weights for all animals across treatment groups. Data are shown as mean ± SD.

**S19:**
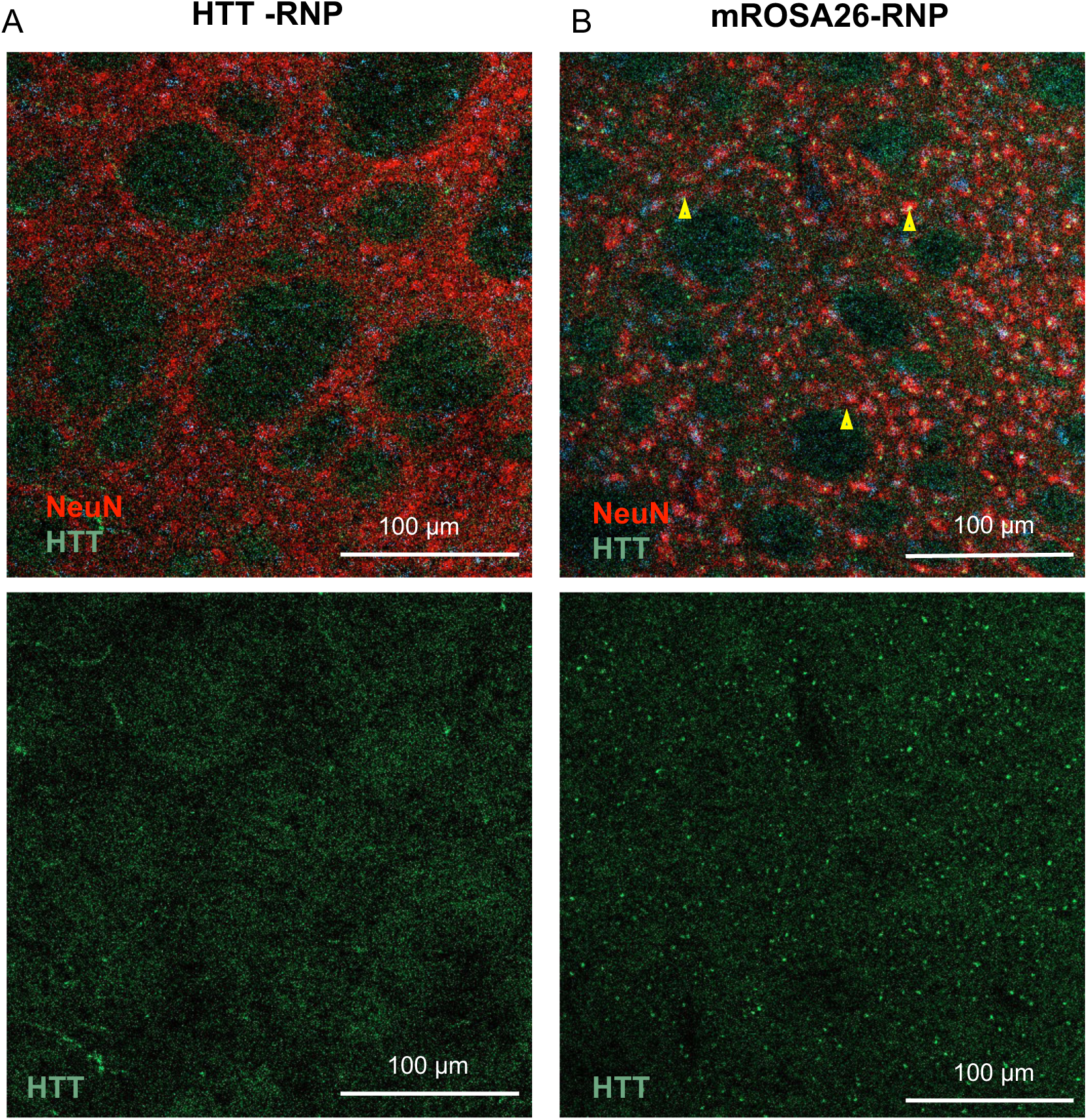
Immunohistochemical detection of mutant huntingtin (mHTT) in striatal sections from R6/2 mice treated with A5K-based Neuro-PERC (A-B) Representative confocal images of striatal tissue from mice treated with HTT-RNP (A) or mROSA26-RNP (B) delivered via A5K-based Neuro-PERC. Sections were stained for NeuN (red) and huntingtin (HTT, green). In the HTT -RNP–treated hemisphere (A), reduced HTT signal was observed relative to the mROSA26 control (B), where persistent mutant HTT aggregates (yellow arrowheads) are evident within NeuN-positive neurons. Scale bars, 100 µm.

**S20:**
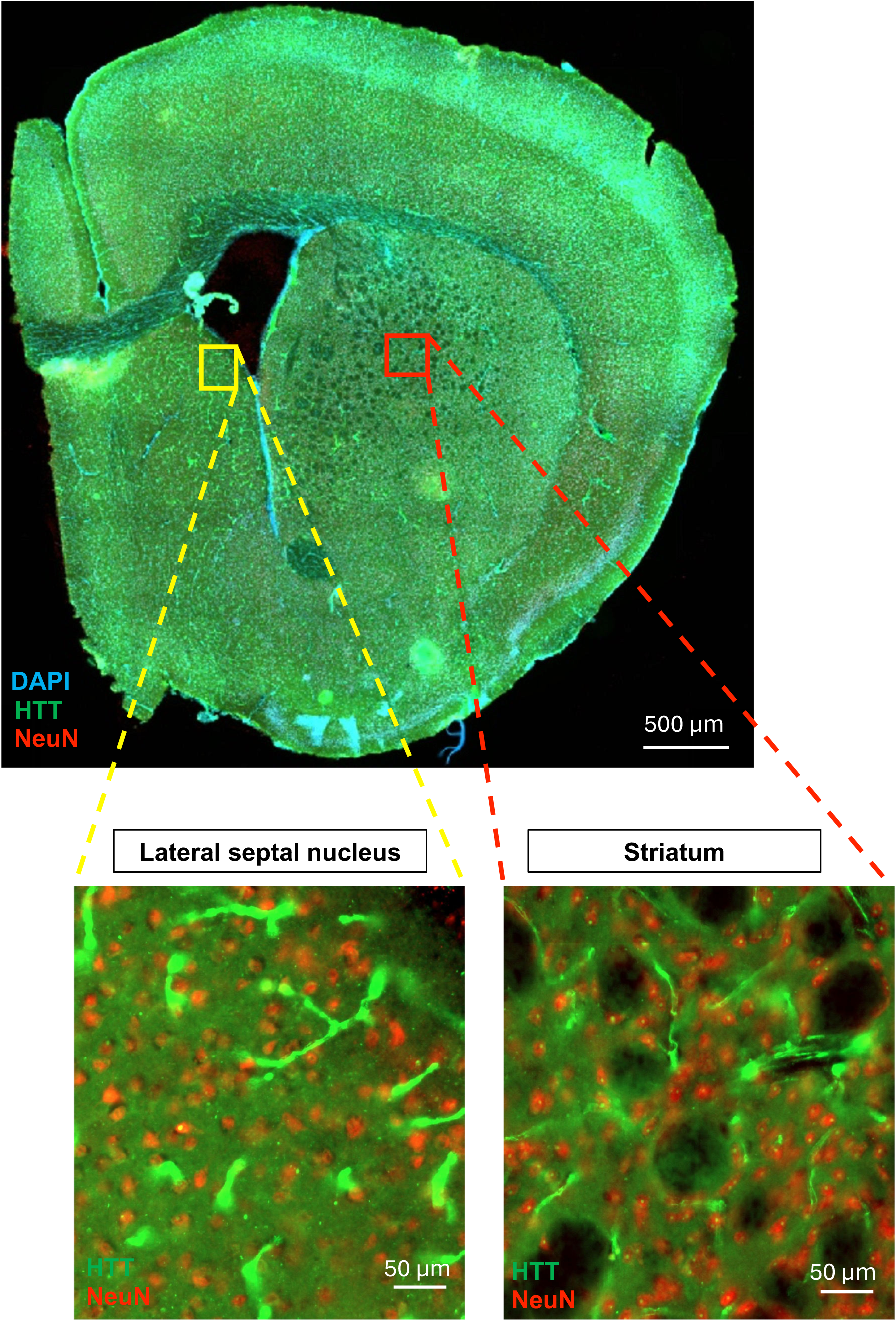
Control mROSA26-RNP–treated mouse brain showing huntingtin (HTT) distribution. Representative confocal image of a coronal brain section from an mROSA26-RNP–treated mouse showing NeuN (red), HTT (green), and DAPI (blue) staining. Insets highlight higher-magnification views of the lateral septal nucleus (yellow box) and striatum (red box). Robust HTT signal is evident in both regions, serving as an internal control for comparison with HTT-targeted RNP treated hemispheres (Fig 4E) Scale bars, 500 µm (whole section) and 50 µm (insets).

## Supplemental tables

**Table S1:**
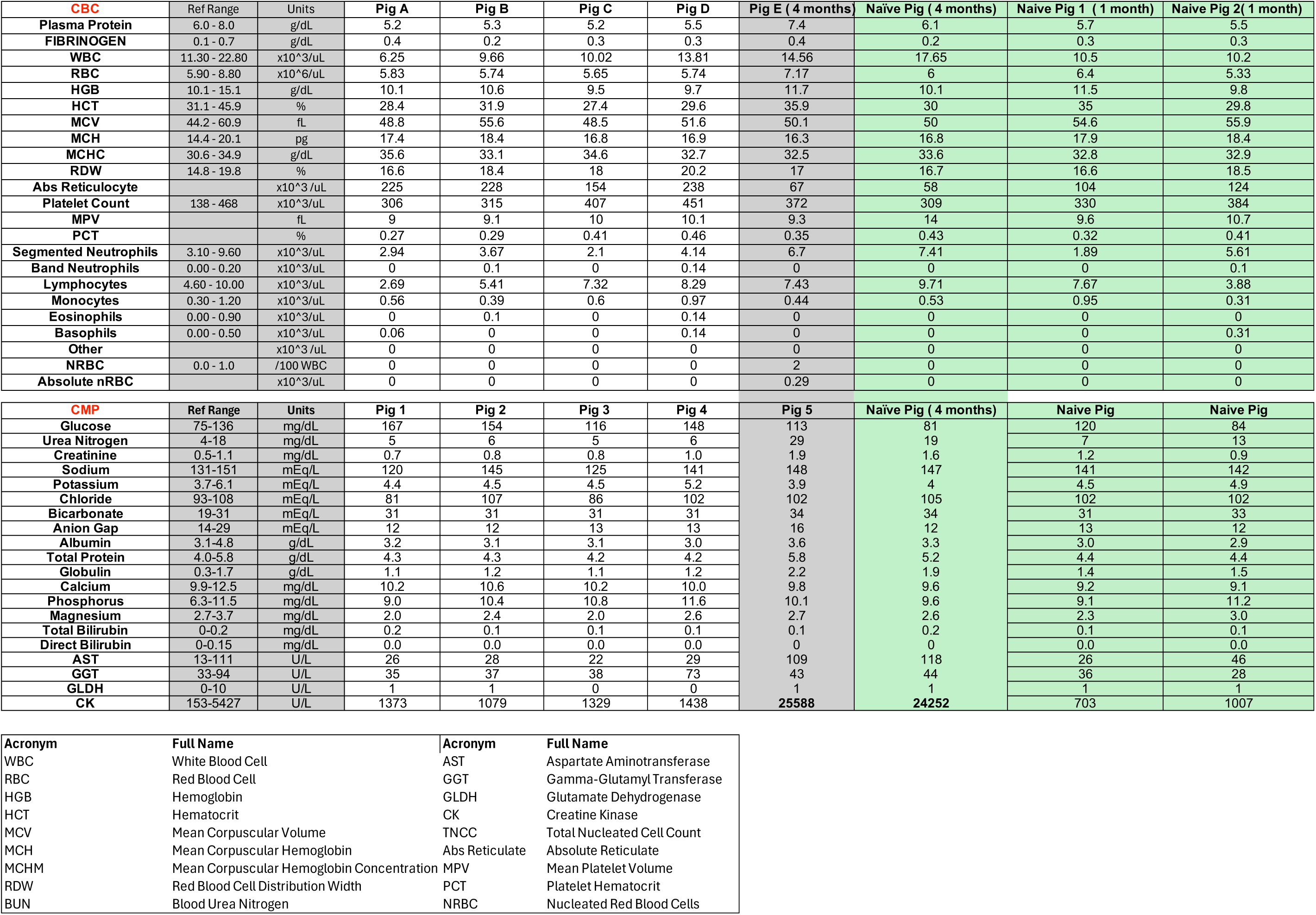
Blood panel from pig experiment. Comprehensive blood analysis of five Neuro-PERC–treated pigs and age-matched naïve controls was performed to assess systemic safety. Complete blood count (CBC; top panel) and clinical metabolic profile (CMP; bottom panel) values are shown alongside reference ranges. Measurements were obtained at 1 month (pigs A-B; naïve age matched pigs: naïve pig 1 (1 month) and naïve pig 2 (1 month)) and at 4 months (pig E; naïve age matched pigs: naïve pig 4 months). Parameters include markers of hematologic status (e.g., RBC, HGB, WBC, platelets), liver function (ALT, AST, GGT, GLDH), renal function (urea, creatinine, BUN), and electrolytes. Reference ranges and acronyms are provided.

**Table S2:**
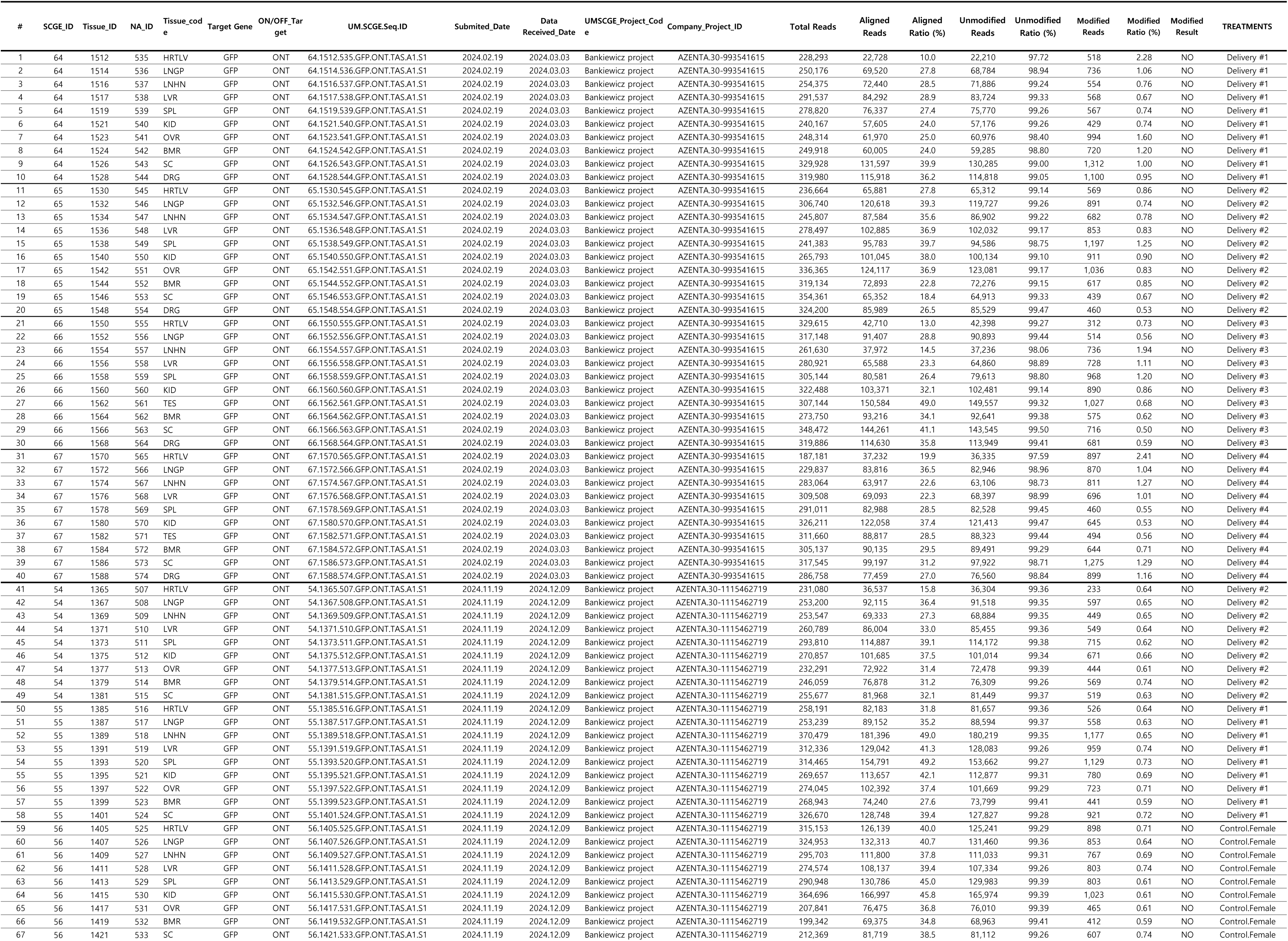
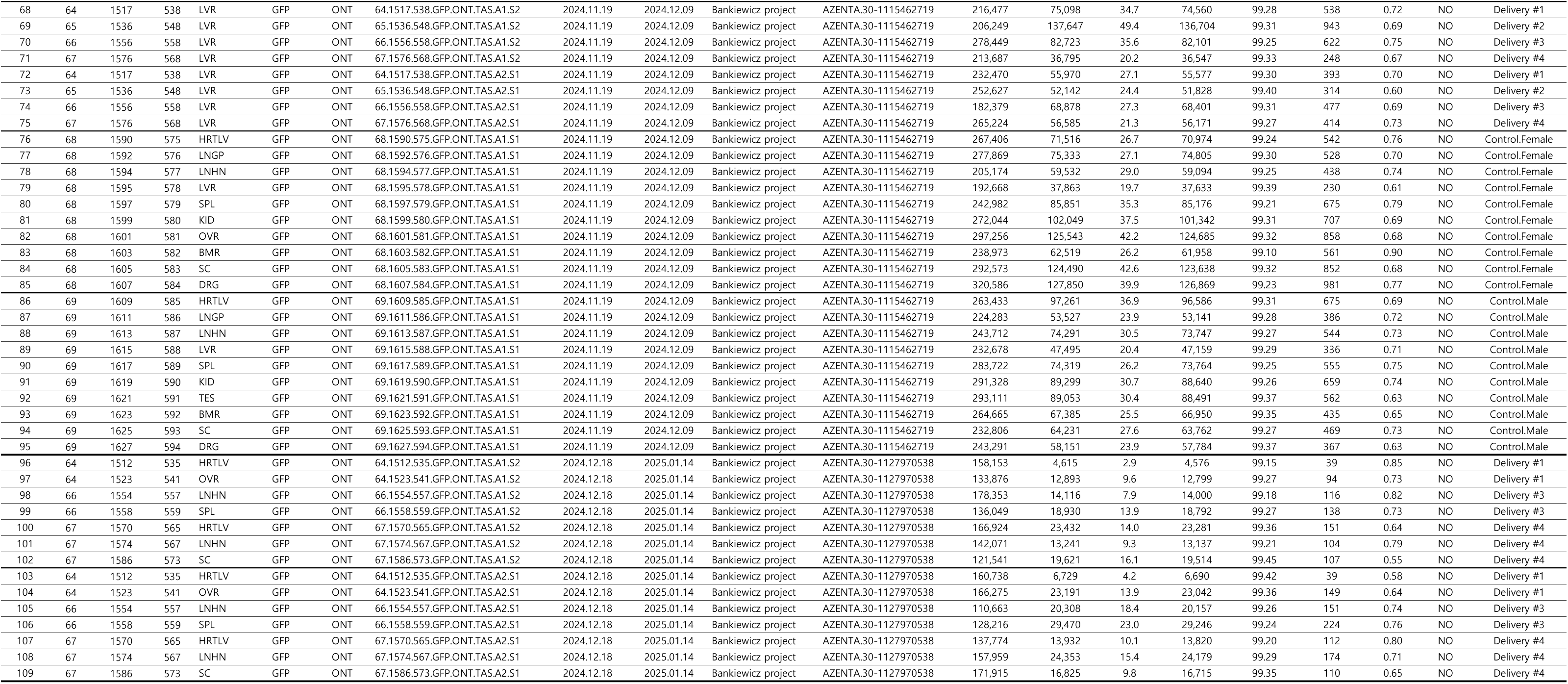
Genome Safety Test using Targeted Amplicon Next Generation Sequencing on pig tissue. Genomic DNA was extracted from peripheral tissues—including heart (HRTLV), lung (LNGP), lymph node (LNHN), liver (LVR), spleen (SPL), kidney (KID), gonads (TES/ OVR), bone marrow (BMR), spinal cord (SC), and dorsal root ganglion (DRG, to assess systemic genome editing following intrastriatal Neuro-PERC administration. Targeted PCR amplification of the GFP locus was performed and amplicons were sequenced using Oxford Nanopore (ONT) long-read sequencing. CRISPResso2 analysis confirmed absence of detectable off-target genome editing. Treated animals corresponded to Delivery Groups A-D (Pig SCGE IDs (column 2): 64, 65, 66, and 67), while Pig 54 and Pig 55 represented additional delivery replicates. Control animals included female controls (Pig 56 and Pig 68) and a male control (Pig 69)

**Table S3:**
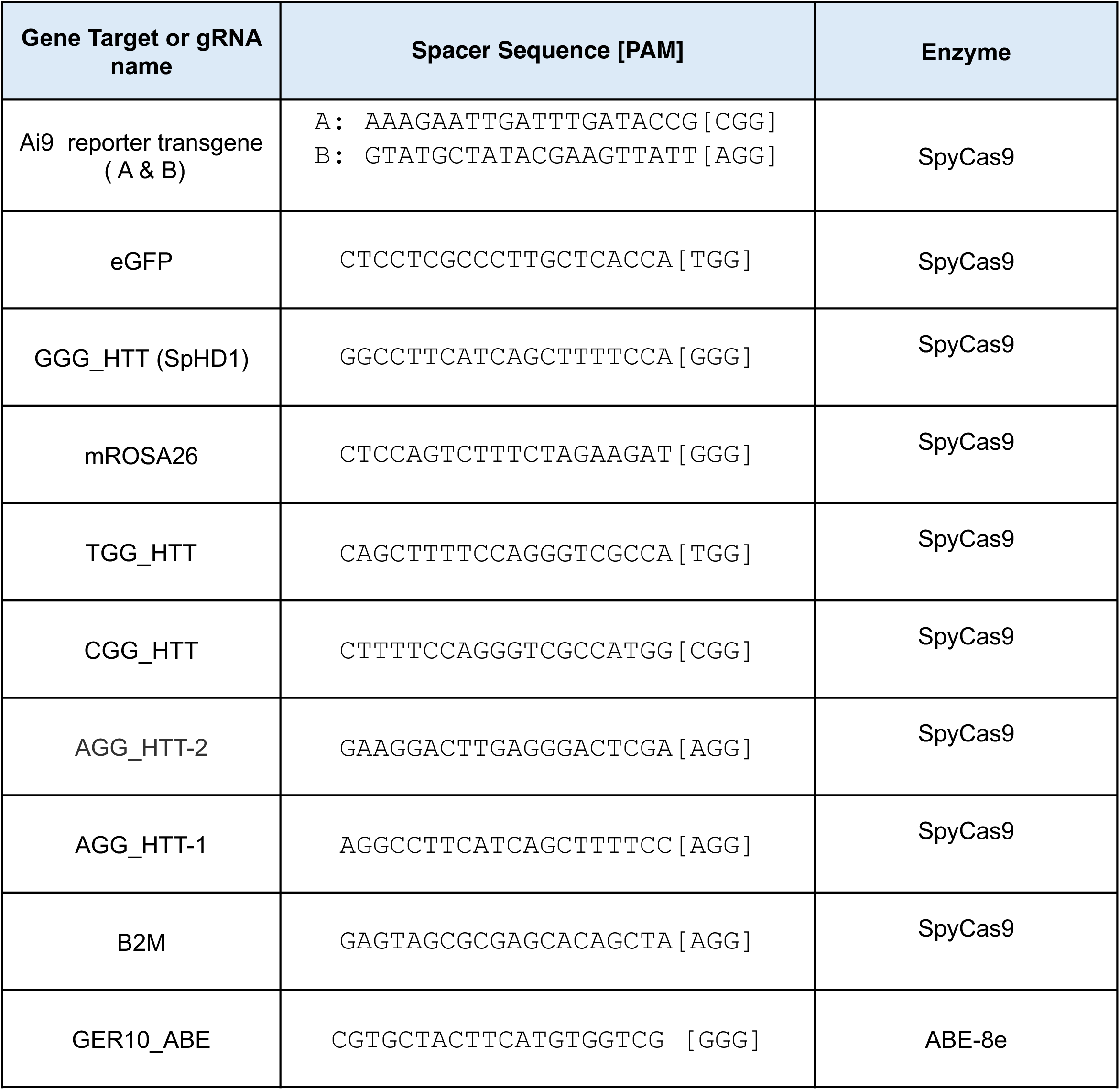
Guide RNAs used.

**Table S4:**
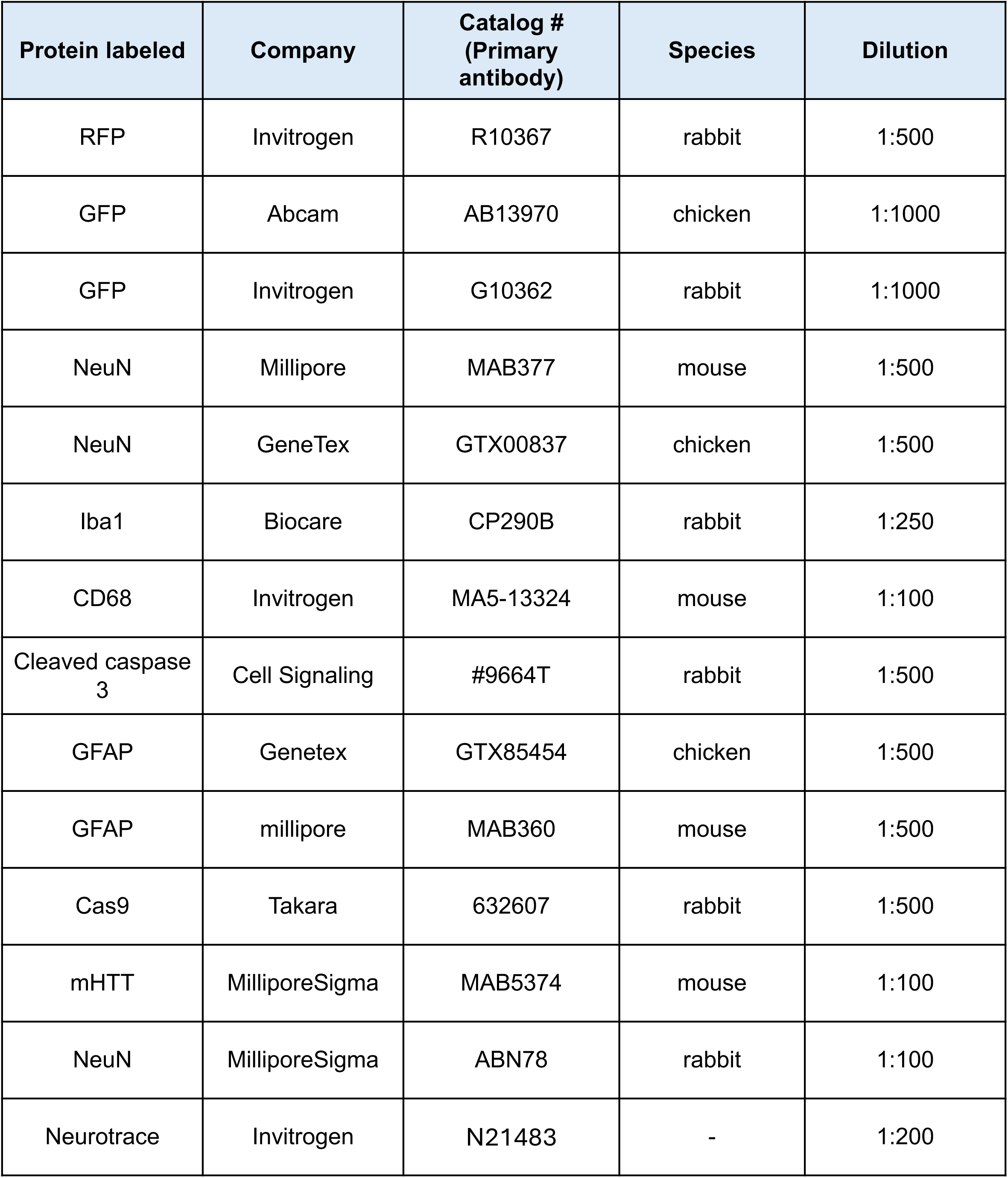
Primary antibodies.

**Table S5:**
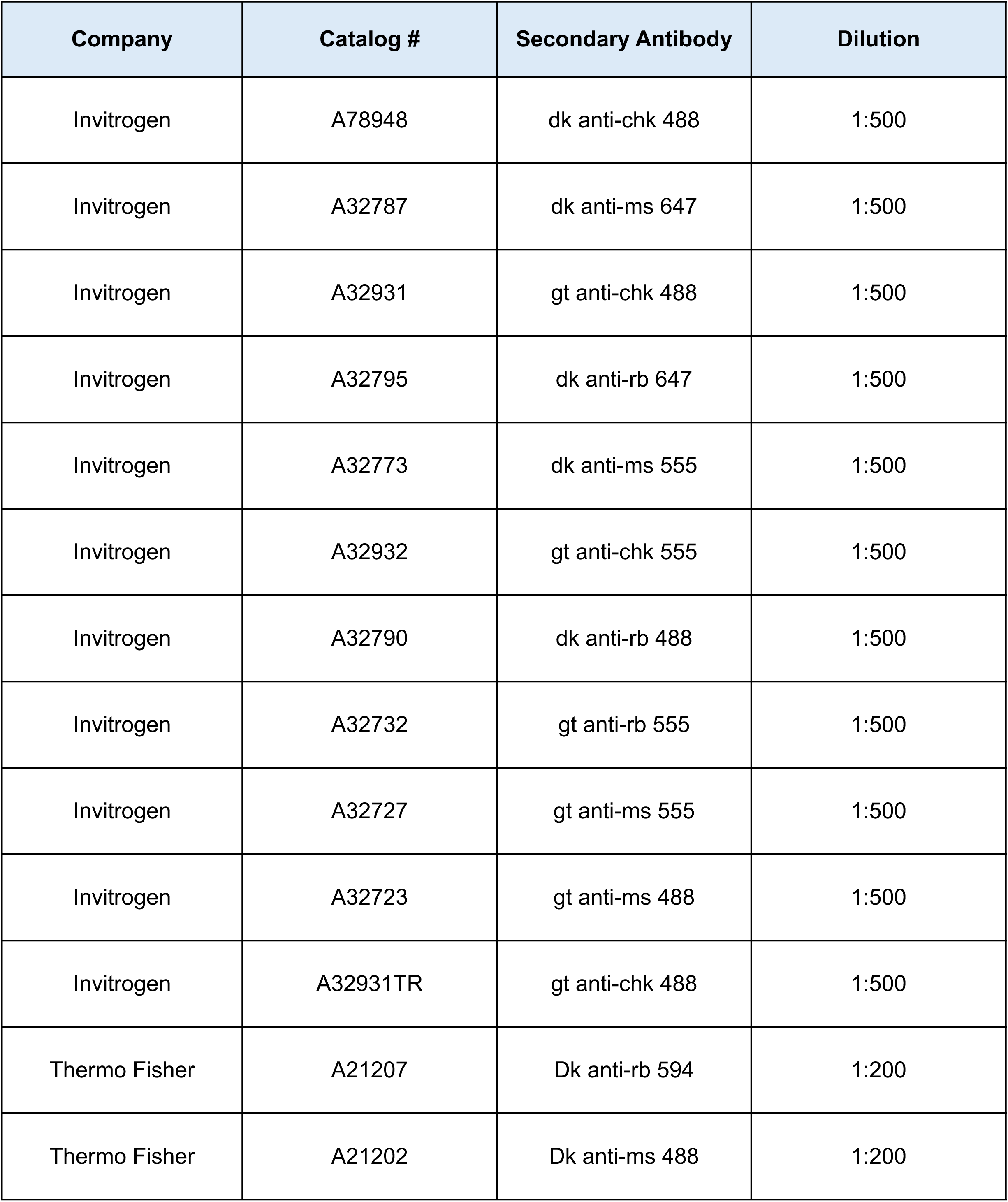
Secondary antibodies.

**Table S6:**
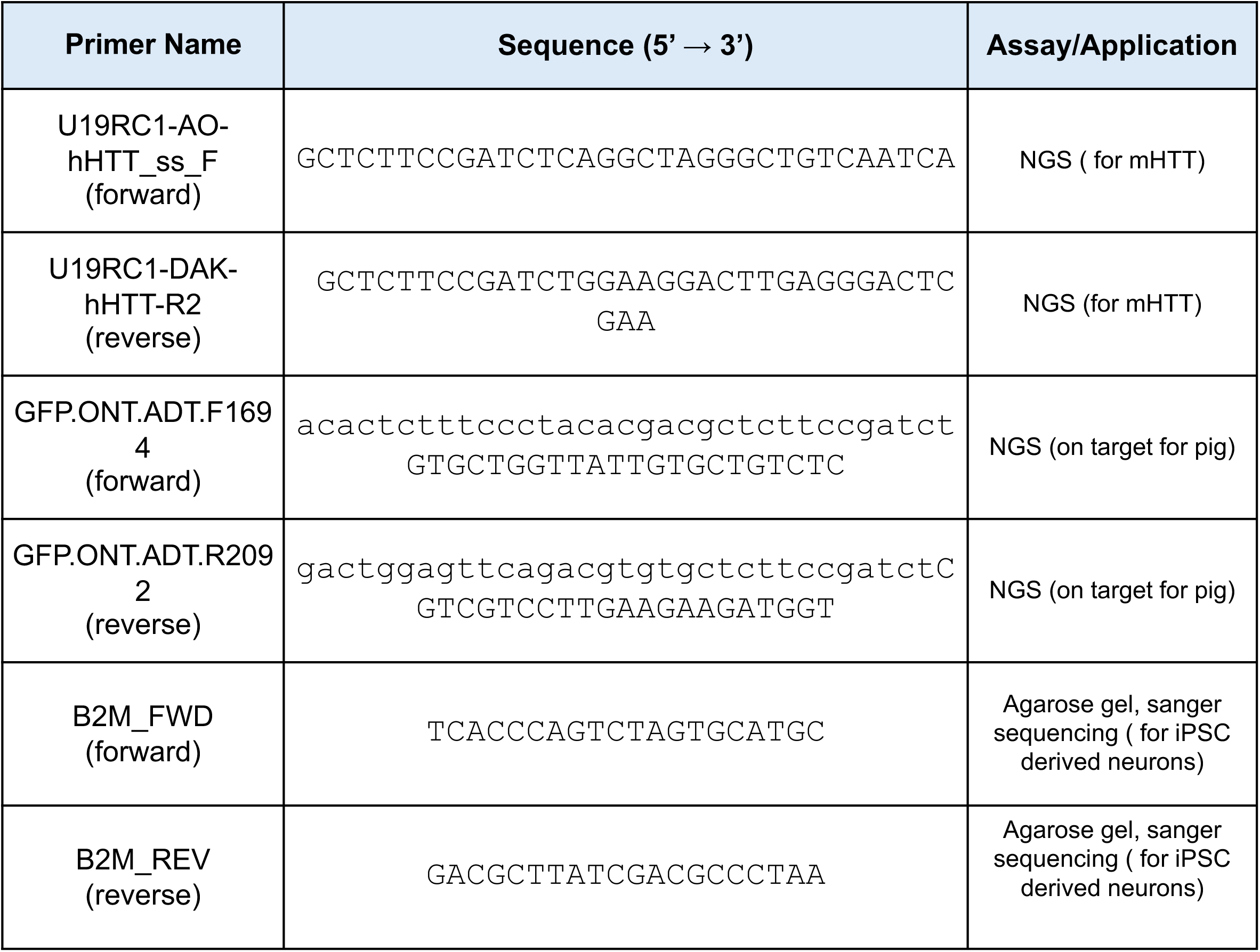
Oligonucleotides for generating amplicons that were analyzed by NGS, sanger sequencing or agarose gels.

